# Dynamic modeling of *Streptococcus pneumoniae* competence provides regulatory mechanistic insights

**DOI:** 10.1101/300814

**Authors:** Mathias Weyder, Marc Prudhomme, Mathieu Bergé, Patrice Polard, Gwennaele Fichant

## Abstract

In the human pathogen *Streptococcus pneumoniae*, the gene regulatory circuit leading to the transient state of competence for natural transformation is based on production of an auto-inducer that activates a positive feedback loop. About one hundred genes are activated in two successive waves linked by a central alternative sigma factor ComX. This mechanism appears to be fundamental to the biological fitness of *S. pneumoniae.* We have developed a knowledge-based model of the competence cycle that describes average cell behavior. It reveals that the expression rates of the two competence operon, *comAB* and *comCDE*, involved in the positive feedback loop must be coordinated to elicit spontaneous competence. Simulations revealed the requirement for an unknown late *com* gene product that shuts of competence by impairing ComX activity. Further simulations led to the predictions that the membrane protein ComD bound to CSP reacts directly to pH change of the medium and that blindness to CSP during the post-competence phase is controlled by late DprA protein. Both predictions were confirmed experimentally.

## Introduction

In *Streptococcus pneumoniae*, competence or X state, is a transient physiological state induced by activation of a genetic program in response to specific conditions, such as environmental stress (Claverys *et al*, 2006). Competence state, at least, allows natural transformation, fratricide (Claverys *et al*, 2007), biofilm formation (Oggioni *et al*, 2006; Trappetti *et al*, 2011; Vidal *et al*, 2013) and contributes to virulence efficiency (Zhu *et al*, 2015; Lin *et al*, 2016). In laboratory exponential cultures, after addition of the synthetic auto-inducer (the competence-stimulating peptide (CSP) (Håvarstein *et al*, 1995), competence develops abruptly and nearly simultaneously in virtually all the cells, and then, after about 20 minutes, declines almost as quickly (Alloing *et al*, 1998). During this short period, the bacteria are able to lyse neighboring cells of their siblings or close relatives (Claverys *et al*, 2007; Johnsborg & Håvarstein, 2009) and to transform their genome by taking up exogenous DNA and incorporating it through RecA-mediated homologous recombination (Martin *et al*, 1995; Chen & Dubnau, 2004; Johnston *et al*, 2014a). This transformation process is natural to many bacterial taxa and is thought to be a driver of evolution through promotion of horizontal gene transfer (Johnston *et al*, 2014b). By facilitating acquisition of new genetic traits, competence thus enables *S. pneumoniae*, a major human pathogen, to adapt to changing environmental conditions by, for example, promoting antibiotic resistance (Tomasz, 1997) and vaccine evasion (Croucher *et al*, 2011; Golubchik *et al*, 2012).

Competence in *S. pneumoniae* results from successive waves transcription of two groups of *com* genes, termed early and late (Dagkessamanskaia *et al*, 2004; Peterson *et al*, 2004) (Figure 1A). Competence develops in response to the export and accumulation of CSP by the membrane ComAB transporter after maturation of the pre-CSP encoded by *comC* (Claverys *et al*, 2006). At a critical concentration, CSP activates the two-component signal transduction system ComDE (Pestova *et al*, 1996). The membrane-bound histidine kinase ComD associated with CSP transmits the signal to its cognate response regulator ComE by a phosphorelay (Martin *et al*, 2013). ComE∼P directly activates the early *com* genes by binding to direct repeats (ComE-box) in the promoters of their operons (Martin *et al*, 2013; Boudes *et al*, 2014). These include the *comAB* and *comCDE* operons, creating a positive feedback loop which amplifies the signal and allows competence propagation throughout the population. Included in the early genes activated by ComE∼P is the central competence regulator gene *comX* which encodes ComX, the competence-specific σ factor (σ^X^) (Lee & Morrison, 1999). ComX enables RNA polymerase to recognize a specific 8 bp sequence (combox or Cinbox) that characterizes the promoters of late *com* genes (Claverys & Havarstein, 2002; Peterson *et al*, 2000). Another early gene, *comW*, (Luo *et al*, 2004) is involved in stabilization of ComX through prevention of ClpP-dependent proteolysis (Piotrowski *et al*, 2009), as well as in ComX-mediated activation (Sung & Morrison, 2005), possibly through enhancement of ComX’s binding to core RNA polymerase (Tovpeko & Morrison, 2014; Tovpeko *et al*, 2016). However, the mechanisms by which ComW acts are still unclear. Genes under direct ComX control include those coding for the DNA uptake machinery and proteins dedicated to the processing of transforming DNA (Johnston *et al*, 2014b), such as DprA, as well as genes implicated in fratricide (Claverys *et al*, 2007). Shutoff of pneumococcal competence depends on two known mechanisms: i) the balance between ComE∼P, which activates early competence genes, and ComE, which antagonizes their expression by competing for binding to the ComE-box (Martin *et al*, 2013), and ii) the repression of the positive feedback loop by DprA, which forms a complex with ComE∼P that blocks the latter’s action through either sequestration or dephosphorylation (Mirouze *et al*, 2013; Weng *et al*, 2013).

**Figure 1.**
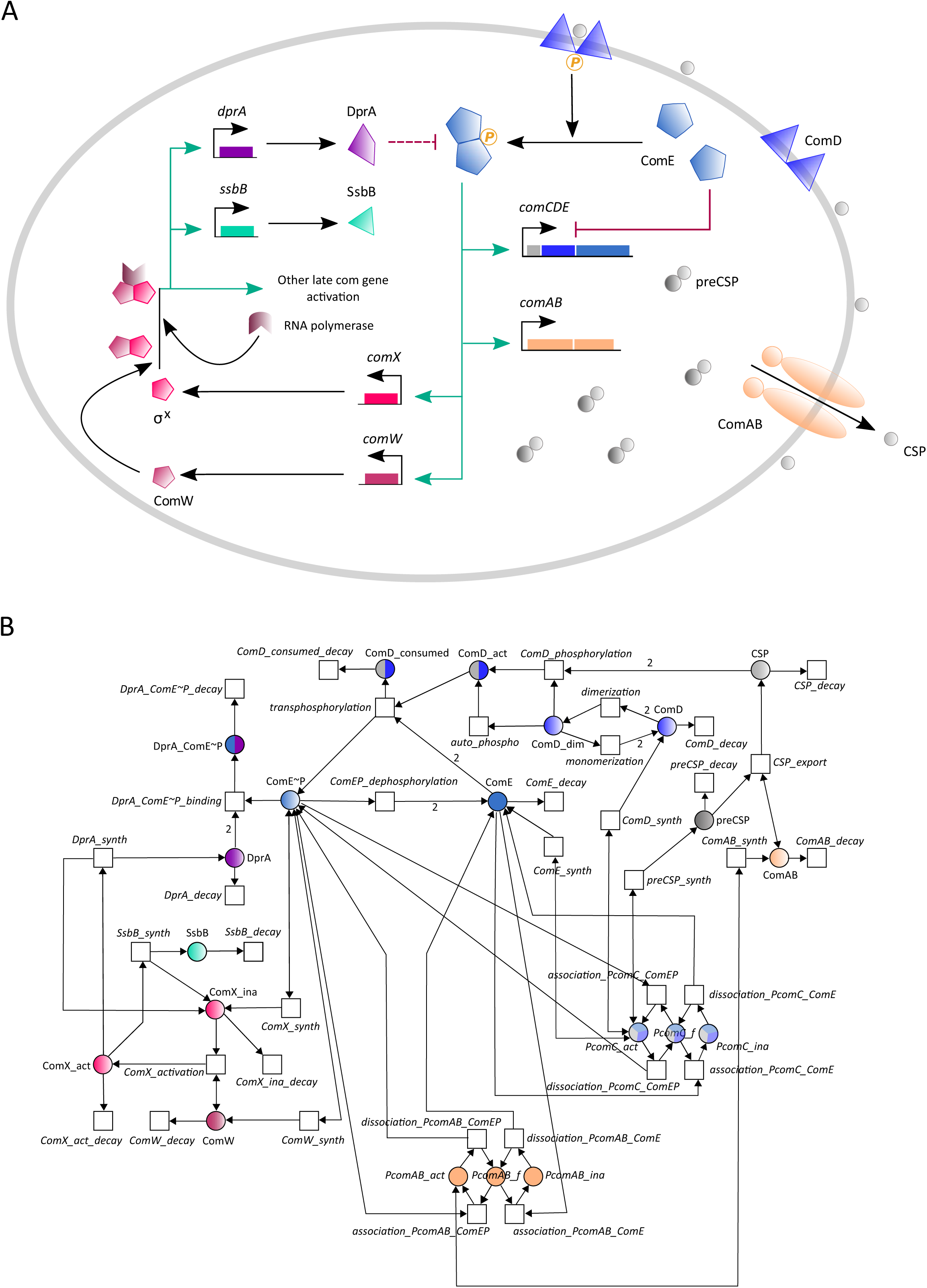
Overview of the competence regulatory network in *S. pneumonia* and its Petri net modeling.

(A) Competence develops in response to a competence-stimulating peptide (CSP) which is first synthesized as a precursor form (pre-CSP) encoded by *comC*. The pre-CSP is matured and exported by the dedicated ABC transporter ComAB. Extracellular CSP binds the histidine kinase ComD and triggers autophosphorylation. ComD∼P activates its cognate response regulator ComE by transferring its phosphoryl group. ComE∼P activates the transcription of early *com* genes which include the *comCDE* and *comAB* operons, establishing a positive feedback loop. Early *com* genes also comprise *comX* and *comW.* comX encodes σ^X^, the competence-specific σ factor that requires ComW for improving its binding to core RNA polymerase. Since the mechanisms that underlie the action of ComW are still unknown, we simplify the scheme by considering that transcription is ensured by a complex composed of σ^X^/ComW/RNA polymerase. σ^X^ controls the activation of large set of late *com* genes represented here by *ssbB*, which is commonly used in experimental assays as reporter gene for late gene expression, and *dprA* whose gene product is involved in competence shut-off by sequestering ComE∼P. Competence shut-off also involves ComE, which inhibits *comCDE* transcription by outcompeting ComE∼P for binding to P_*comC*_. The green arrows represent activation of gene expression, the red lines represent inhibition reactions and the black arrows represent other reactions like synthesis, binding or export reactions. (B) Petri net model. Circles correspond to places and represent proteins involved in the system. Squares correspond to transitions and represent the system reactions. Places and transitions are connected by directed weighted arcs (arrows) whose associated number corresponds to the stoichiometric coefficient of the reaction when it differs from one. Places are colored according to the color code adopted in (A). Names of places and transitions are indicated. Bidirectional arrows represent test arcs.

The circuits that regulate competence for transformation are adapted to the lifestyle of each species, as exemplified by *S. pneumoniae* and *B. subtilis* (Johnston *et al*, 2014b; Claverys *et al*, 2006). Two distinct regulatory circuits have been reported to control the expression of ComX within the Streptococci. Phylogenetic analyses have shown that one, based on the *comCDE* system, is present in species of the mitis and anginosus groups (Martin *et al*, 2006) while the other, based on the recently-discovered *comRS* transcriptional activation system (Gardan *et al*, 2009; Fontaine *et al*, 2010), is found in species of the mutans, salivarius, bovis, pyogenic and suis groups (Johnston *et al*, 2014b; Fontaine *et al*, 2015).

Several mathematical models have been developed to aid the study of competence regulation in *B. subtilis* (Maamar & Dubnau, 2005; Süel *et al*, 2006; Maamar *et al*, 2007; Schultz *et al*, 2007; Leisner *et al*, 2009; Schultz *et al*, 2009, 2013). Two models have also been published to help answer some of outstanding questions concerning the ComRS regulatory cascades of *S. mutans* (Son *et al*, 2012) and *S. thermophiles* (Haustenne *et al*, 2015). Both models were established to investigate components of the ComRS system critical for ComX production. However, despite our extensive knowledge of ComCDE regulation circuit in *S.pneumoniae*, only two attempts to model this regulatory circuit have been published. Karlsson and collaborators focused on a possible mechanism for abrupt competence shut-off, since the mechanisms involved had not yet been unraveled (Karlsson *et al*, 2007). Their model suggested that a putative *comX*-dependent repressor that inhibits expression of *comCDE* and *comX*, is responsible for competence shut-down. However, subsequent work argued against this proposal, by showing that the late competence protein DprA is involved in competence shutoff through dephosphorylation or sequestration of ComE∼P (Mirouze *et al*, 2013) rather than through direct interference with *comCDE* and *comX*expression. More recently, Moreno-Gámez and collaborators (Moreno-Gámez *et al*, 2017) published a model of spontaneous competence development in *S. pneumoniae* that takes account of environmental conditions and cell history. Both models describe competence development for a homogeneous population, based on the assumption that initiation of competence is controlled by a quorum sensing system in which cell density must rise high enough for the population to detect the CSP auto-inducer threshold and switch en masse to the competent state.

However, other observations challenge the view that CSP accumulation in the growth medium is directly proportional to the cell density. Experiments have shown that various environmental parameters control the timing of spontaneous competence induction in cell populations, disqualifying cell density as the crucial parameter (Claverys *et al*, 2006). Recent results (Prudhomme *et al*, 2016) show that the spontaneous competence shift in a non-competent population relies on a self-activated cell sub-population that arises *via* a growth time-dependent mechanism. During this short period of competence development, since CSP is mostly retained on the cell surface, competence propagates by successive contacts between activated and non-competent receiver cells. These results call into question the assumption of population homogeneity that underlies the former mathematical models of competence regulation in *S. pneumoniae.* Indeed, designing a dynamic model at the population scale requires one to take into account non-homogeneity of the population and interaction between individual cells. We have now modelled this regulatory circuit, first at the cell level to provide a module that can then be embedded in more complex models to study competence propagation within the whole cell population where both hypotheses - quorum sensing or self-activated sub-population - can be tested.

We have taken a two-step approach. We first integrated the available information into a Petri net framework and performed structural analysis of the model. We then explored the dynamics of the model in a deterministic framework, using ordinary differential equations (ODEs). We have exploited previously published real time measurements of gene activities obtained from *in vivo* transcriptional data (Mirouze *et al*, 2013), which we have transformed into average promoter activities and average protein synthesis rates per cell by applying recently published mathematical approaches (de Jong *et al*, 2010; Stefan *et al*, 2015). The first designed knowledge-based model corroborates previous experimental findings but also reveals gaps in our knowledge of competence shut-off. We have addressed these gaps by testing eight alternative models for competence shut-off. Our results suggest that competence shut-off involves an interaction between ComW and the product of a late *com* gene which impairs ComX activity. Furthermore, *in silico* perturbations of key network parameters revealed the mechanism of two hallmarks of pneumococcal competence regulation: i) modulation of competence induction level by pH variation in the growth medium appears to be directly linked to CSP interaction with ComD; ii) cells exiting competence could not immediately re-start a competence cycle, a ‘blind-to-CSP’ period found to be controlled by DprA. Both were supported experimentally. Moreover, using the model to simulate spontaneous competence induction, we highlight the need for coordinated basal expression of *comAB* and *comCDE* to govern the ComE∼P-ComE ratio that is crucial to initiating the positive feedback loop.

## Results

### Modeling assumptions and reaction network description

We focused our efforts on creating a model whose simulated behaviors are consistent with the available experimental results. Below, we give more details on the available knowledge concerning some important parts of the regulatory network, in order to explain our modeling choices and the assumptions we made to simplify our model. The reactions are summarized in Table 1 and depicted in Figure 1A. We addressed the dynamic modeling of the *S. pneumoniae* competence cycle at the cell level, where competence is induced in laboratory conditions by addition of saturating concentrations of exogenous synthetic CSP to non-competent cells growing in liquid culture. Induction in this manner triggers competence synchronously throughout the population. Hence, quantitative measurements of reporter gene expression can be used to estimate the average cell behavior (de Jong *et al*, 2010; Stefan *et al*, 2015).

**Table 1.** Reactions involved in competence regulation

| Description | Reaction number | Reaction | Petri net transition |
| --- | --- | --- | --- |
| *Maturation and export of comC product (pre-CSP) by the dedicated ABC transporter ComAB* | 1 | $\text{pre-CSP} + \text{ComAB} \rightarrow \text{CSP} + \text{ComAB}$ | CSP_export |
| *Dimerization of the histidine kinase ComD* | 2 | $2 \text{ ComD} \rightarrow (\text{ComD})_D$ | dimerization |
| *Dissociation of $(\text{ComD})_D$* | 3 | $(\text{ComD})_D \rightarrow 2 \text{ ComD}$ | monomerization |
| *Basal autophosphorylation of $(\text{ComD})_D$* | 4 | $(\text{ComD})_D \text{ phosphates} \rightarrow (\text{ComD}_{\text{act}})_D$ | auto_phospho |
| *CSP-induced autophosphorylation of $(\text{ComD})_D$* | 5 | $2 \text{ CSP} + (\text{ComD})_D + \text{phosphates} \rightarrow (\text{ComD}_{\text{act}})_D$ | ComD_phosphorylation |
| *Activation of ComE as a dimer by phosphoryl group transfer from $(\text{ComD}_{\text{act}})_D$* | 6 | $(\text{ComD}_{\text{act}})_D + 2 \text{ ComE} \rightarrow (\text{ComE}\sim\text{P})_D + (\text{ComD}\text{-CSP})_D$ | transphosphorylation |
| *Self-catalyzed dephosphorylation of $(\text{ComE}\sim\text{P})_D$* | 7 | $(\text{ComE}\sim\text{P})_D \rightarrow 2 \text{ ComE} + \text{phosphates}$ | ComEP_dephosphorylation |
| *Early com gene product synthesis through activation of their genes by $(\text{ComE}\sim\text{P})_D$* | 8 | $(\text{ComE}\sim\text{P})_D \rightarrow \text{ComE} + \text{ComD} + \text{ComAB} + \text{pre-CSP} + \text{ComW} + \text{ComX}_{\text{ina}} + (\text{ComE}\sim\text{P})_D$ | ComD_synth; ComE_synt;<br>ComAB_synth;<br>preCSP_synth;<br>ComW_synth;<br>ComX_synth |
| *ComX activation* | 9 | $\text{ComX}_{\text{ina}} + \text{ComW} \rightarrow \text{ComX}_{\text{act}} + \text{ComW}$ | ComX_activation |
| *Late com gene product synthesis through activation of their genes by the active sigma factor ComX* | 10 | $\text{ComX}_{\text{act}} \rightarrow \text{ComX}_{\text{ina}} + \text{DprA} + \text{SsbB}$ | DprA_synth; SsbB_synth |
| *Inactivation of $\text{ComE}\sim\text{P}$ by sequestration in a complex with DprA* | 11 | $2 \text{ DprA} + (\text{ComE}\sim\text{P})_D \rightarrow (\text{DprA}\text{-}\text{ComE}\sim\text{P})_D$ | DprA_ComEP_binding |
| *Degradation of network components* | 12 | $\begin{aligned} &\text{pre-CSP} \rightarrow \emptyset; \text{ComAB} \rightarrow \emptyset; \\ &\text{CSP} \rightarrow \emptyset; \text{ComD} \rightarrow \emptyset; \\ &(\text{ComD}\text{-CSP})_D \rightarrow \emptyset; \text{ComE} \rightarrow \emptyset; \\ &\text{ComX}_{\text{act}} \rightarrow \emptyset; \text{ComX}_{\text{ina}} \rightarrow \emptyset; \\ &\text{ComW} \rightarrow \emptyset; \text{DprA} \rightarrow \emptyset; \\ &\text{SsbB} \rightarrow \emptyset \\ &(\text{DprA}\text{-}\text{ComE}\sim\text{P})_D \rightarrow \emptyset; \end{aligned}$ | $\begin{aligned} &\text{preCSP\_decay}; \text{ComAB\_} \\ &\text{decay}; \text{CSP\_decay}; \\ &\text{ComD\_decay}; \\ &\text{ComD\_consumed\_decay}; \\ &\text{ComE\_decay}; \\ &\text{ComX\_act\_decay}; \\ &\text{ComXina\_decay}; \\ &\text{ComW\_decay}; \\ &\text{DprA\_decay}; \text{SsbB\_decay} \\ &\text{DprA}\text{-}\text{ComE}\sim\text{P\_decay}; \end{aligned}$ |

### Assumptions

To simplify analysis and computational handling, we treated transcription and translation as a single step, and designed our network at the protein level. This simplification is based on the assumption that the mRNA concentrations are in quasi-steady state, in the sense that they adapt almost instantaneously to changes in promoter activity. Thus, it is possible to overlook the variations of mRNA concentration and to write variations of the protein concentration directly as a function of the promoter activity. This is known as the quasi-steady-state approximation. Indeed, when slow processes dominate, the fast processes are assumed to be continuously in quasi-equilibrium (Chen *et al*, 2010).

Further assumptions were made to reduce the number of actors in the network : i) CSP degradation by the cell wall protease HtrA (Cassone *et al*, 2012) was not explicitly modeled but was included in the CSP degradation constant; ii) ATP and ADP cofactors for phosphorylation reactions were not explicitly included in the network, since ATP concentration is presumably not a limiting factor during exponential growth; iii) among the late *com* gene products, *i.e.* products of genes under control of ComX, we took into account only DprA, whose action in competence shut-off has been described (Mirouze *et al*, 2013), and SsbB, which is commonly used in experimental assays as a reporter of late gene expression; iv) ComX, being a sigma factor, regulates its target genes by forming a complex with RNA polymerase, but since only *comX* expression is regulated by the competence network we have not explicitly included the RNA polymerase component; v) DprA-ComE binding was ignored, since there are no quantitative data on DprA-ComE∼P protein-protein interactions, other than that DprA has much higher affinity for ComE∼P than for ComE (Mirouze *et al*, 2013).

#### Initiation of the competence state: the ComDE two-component system

Once CSP reaches a critical external concentration, it activates the two-component signal-transduction system (TCS) ComDE, which belongs to the AgrA/AlgR/LytR family (Lange *et al*, 1999). Since the ComDE mechanism is not completely understood, we took advantage of the experimental results obtained from detailed studies performed in *Staphylococcus aureus* on the AgrCA TCS (George Cisar *et al*, 2009). The AgrC histidine kinase forms a dimer before autophosphorylation. This dimer possesses a basal autophosphorylation activity independent of ligand binding, as also observed for ComD (Martin *et al*, 2010). The AgrC dimer possesses two independent ligand binding sites with no evidence of cooperativity (Wang *et al*, 2014). By analogy, we have modelled ComD phosphorylation in the same manner: formation of a dimer of ComD (Table 1, reaction 2 and reverse reaction 3) whose autophosphorylation can be either independent of CSP binding (Table 1, Reaction 4) or activated by CSP interaction (Table 1, reaction 5).

Despite several attempts, transphosphorylation of ComE *in vivo* to the active form that induces early genes was not detected (Martin *et al*, 2013). However, the phosphorylmimetic mutant ComE^D58E^ protein dimerizes in solution whereas ComE has been observed only in its monomeric form (Martin *et al*, 2013). Moreover, structural data on the complex ComD/ComE/*comCDE* indicates that the transfer of the phosphoryl group from the histidine kinase to its cognate response regulator ComE mediates ComE dimerization through the binding of a phosphorylated ComD dimer to two monomers of ComE (Sanchez *et al*, 2015). Hence, we simplify the reaction by assuming that a dimer of ComD∼P transphosphorylates a dimer of ComE (Table 1, reaction 6). After transphosphorylation, the CSP-bound ComD dimer ((ComD-CSP)_D_) can either enter a cycle of phosphorylation-transphosphorylation and activate many molecules of ComE or become inactive. The fact that the number of ComD molecules increases from 1500 per cell before competence induction to 39000 per competent cell (Martin *et al*, 2013) and the recent demonstration that ComD is involved in CSP retention (Prudhomme *et al*, 2016) tend to favor the second hypothesis. Therefore, we assume that once an active CSP-bound dimer of ComD has phosphorylated a dimer of ComE, it becomes inactive. We simply model the fate of the inactive form of the CSP-bound dimer of ComD by degradation (Table 1, reaction (12)).

The dephosphorylation of ComE∼P has not yet been documented. No phosphatase active on the histidine kinase ComD has been reported, nor has any other protein that might fulfill this function. Moreover, no detectable phosphatase activity of the homologous *S. aureus* AgrC histine kinase on its cognate response regulator AgrA was detected, and it was concluded that the decrease in AgrA phosphorylation level was due to its self-catalyzed dephosphorylation (Wang *et al*, 2014). Consequently, in our model, we hypothesize that ComE∼P will also catalyze its own dephosphorylation (Table 1, reaction 7).

Since the degradation rates of (ComD∼P-CSP)_D_, also called ComD_act_ in Table 1, and of (ComE∼P)_D_are very slow compared to their phosphorylation lability (see below in “ parameter estimation”), we did not include reactions that degrade their phosphorylated forms. We considered that these forms are consumed by the transphosphorylation and the dephosphorylation reactions, respectively (Table 1, reactions 6 and 7).

#### The central regulator ComX

The sigma factor ComX, responsible for activating expression of late *com* genes, required ComW both for protection from degradation by ClpE-ClpP protease (Piotrowski *et al*, 2009) and for its activation (Sung & Morrison, 2005). It was first suggested that competence and late gene transcription might terminate due simply to the disappearance of ComX. However, more recent observations, showing that *clpP* mutant cells, in which ComX and ComW are stable, escape from the competent state as rapidly as wild-type, suggested that another mechanism is responsible for terminating late gene transcription (Piotrowski *et al*, 2009). To integrate this knowledge into our model, we uncouple the two roles of ComW. ComX stabilization has been included in the half-life parameter of ComX to avoid incorporating ClpE-ClpP protease in our model. Activation of ComX by ComW has been taken into account by creating two forms of ComX, inactive and active. ComX is first synthesized as an inactive form that becomes active under the action of ComW (Table 1, reactions 8 and 9).

#### Competence shut-off

It has been recently suggested that the shut-off of the *comCDE* promoter (P_comC_) is intrinsic to ComDE, as the decrease of *comCDE* transcription occurs in the absence of any late *com* gene product (Martin *et al*, 2013; Mirouze *et al*, 2013). Based on the observation that nonphosphorylated ComE efficiently binds P_comC_ *in vitro* and that its overexpression antagonizes spontaneous competence, it has been proposed that ComE accumulating in response to CSP efficiently outcompetes ComE∼P for binding to P_comC_, thus preventing further transcription (Martin *et al*, 2013). However, almost no antagonization of the *comX* gene promoter (P_comX_) by non-phosphorylated ComE was observed, and lower affinity of ComE for this promoter (Martin *et al*, 2013) suggests that ComE does not inhibit P_comX_ efficiently. In fact, shut-off of P_comX_ requires the action of DprA, a late gene product (Mirouze *et al*, 2013; Weng *et al*, 2013). Yeast two-hybrid assays have identified a strong interaction between DprA and ComE∼P, suggesting a genuine physical interaction, and a weaker interaction with ComE (Mirouze *et al*, 2013; Weng *et al*, 2013). It has also been shown that dimerization of DprA is required for the early *com* gene transcription shut-off. Thus, the action of DprA would be to shift the ComE/ComE∼P ratio in favor of early *com* gene promoter repression. The mechanism by which DprA acts on this ratio is not yet known, and two hypotheses have been proposed: DprA forms a complex with ComE∼P that blocks ComE∼P action through ComE∼P sequestration; alternatively, DprA promotes ComE∼P dephosphorylation. We chose to implement the sequestration scenario in our network since DprA-ComE interaction has been shown but there is no evidence that the interaction leads to dephosphorylation ((Table 1, reaction 11). As DprA and ComE share similarly long half-lives (Martin *et al*, 2013; Mirouze *et al*, 2013), we considered that the fate of the complex is degradation of both proteins (Table 1, reaction 12).

### Mathematical modeling of the competence regulatory network

We first developed a qualitative cell-based model using a Petri net formalism to check the structural consistency of our model by taking advantage of its mathematical formalism. Moreover, a Petri net approach, by providing a simple and easily understood qualitative graphical representation of the network structure, facilitates exchanges during the process of network design between biologists and modelers. The dynamic behavior of the network was further studied by turning the network into a set of ordinary differential equations (ODEs).

#### Petri net modeling

In the Petri net (Figure 1B), the molecular species involved in the reactions of Table 1 constitute places and the reactions have been turned into transitions. For most of the reactions, translation into places and transitions is self-explanatory, and the network structure will not be described in detail. The correspondences between reactions and transitions can be found in Table 1. To model the activation and inhibition of the *comCDE* operon, we introduced three additional places corresponding to the free (*PcomC*_*f*), active (*PcomC*_*act*) and inactive (*PcomC*_*ina*) forms of the *comCDE* operon promoter. Binding of (ComE∼P)_D_ switches the promoter to its active form (transition *association*_*PcomC*_*ComEP*). This active form allows the synthesis of pre-CSP, ComD and ComE (transitions *preCSP*_*synth*, *ComD*_*synth* and *ComE*_*synth* respectively). The same scheme was repeated for the activation and inhibition of *comAB* synthesis through the introduction of the same three forms of the *comAB* promoter. During competence shut-off, the competition between ComE and (ComE∼P)_D_ for binding to P_*comC*_ and P_*comAB*_ will first cause the dissociation of (ComE∼P)_D_, that will switch the active promoter to its free form (transitions*dissociation*_*PcomC*_*ComEP*, *dissociation*_*PcomC*_*ComAB*), followed by binding of ComE leading to the promoter inactivation (*PcomC*_*ina*, *PcomAB*_*ina*) and consequently to inhibition of gene expression (transitions*association*_*PcomC*_*ComE*, *association*_*PcomAB*_*ComE*). As ComE and (ComE∼P)_D_ are competed, the reverse transitions were also modeled: starting from an inactive form of P_*comC*_ (P_*comAB*_), the dissociation of ComE results in a free promoter able to bind (ComE∼P)_D_ (transitions *dissociation*_*PcomC*_*ComE* and *association*_*PcomC*_*ComEP*; *dissociation*_*PcomAB*_*ComE* and *association*_*PcomAB*_*ComEP*). For the regulation of *comX* and *comW* expression, as we assume no direct inhibitory effect of ComE, we simplify the model by omitting the promoter. Thus, (ComE∼P)_D_ directly activates the synthesis of ComX and ComW.

#### Qualitative Petri net model validation

Since we will study the dynamics of this network by using ODEs, we simply perform a qualitative validation of our models that depends only on the graph structure and does not require an initial marking. We used the software Charlie (Heiner *et al*, 2015) to compute the structural invariants (T- and P-invariants) that prove the structural consistency of the model. We obtained a set of 19 minimal T-invariants among which eight are non-trivial (Appendix Figure S1). All described T-invariants are biologically meaningful. As every transition of our network participates in a T-invariant, the model is covered by T-invariants. Thus, every reaction in the system may occur as part of the basic behavior of the Petri net (Koch & Heiner, 2008). This is an important property, as the competence state is a transient physiological process. Thus our model allows a cell in its original growth phase to enter transiently into competence and to return to its original state of growth.

A P-invariant corresponds to a set of places assuring mass conservation and avoiding an infinite increase of molecules in the model. Only two invariants are detected in the model that correspond to the different forms (free, inactive or active) of the two promoters P_*comC*_ and P_*ComAB*_. This result was expected since proteins required to set up this specific physiological state need to be synthesized when the cell enters into competence and further degraded when the system has been shut-off to allow the cell to come back to its original growth phase state. As the synthesis transition of each component has been associated with its cognate degradation transition, there is no risk of an infinite accumulation of a given molecule.

The structural consistency of our model being proven, the network was converted into a set of ODEs.

#### Quantitative dynamic modeling

Each place of the Petri net model representing a molecular species will correspond to a state variable *x_i_(t)*. For a given state variable, all the transition arcs pointing towards the place will contribute to the “gain” rate and all the transition arcs pointing away from the place will contribute to the “loss” rate. The kinetics of the 15 molecular entities of our network, corresponding to the 15 places of the Petri net, were expressed as the 15 differential equations reported in Table 2.

**Table 2.** Ordinary differential equations of the initial model

|  |
| --- |
| $\frac{d[preCSP]}{dt} = \beta_{ComCDE} + v\max_{ComCDE} \frac{[(ComE \sim P)_D]}{[(ComE \sim P)_D] + K_{ComE \sim P} * \left(1 + \frac{[ComE]^e}{Ki_{ComE}^e}\right)} - \varepsilon * [ComAB] * [preCSP] - \gamma_{preCSP} * [preCSP]$ |
| $\frac{d[CSP]}{dt} = \varepsilon * [ComAB] * [preCSP] - 2 * v\max_{act} * \frac{[CSP]}{[CSP] + K_{act}} * [(ComD)_D] - \gamma_{CSP} * [CSP]$ |
| $\frac{d[ComAB]}{dt} = \beta_{ComAB} + v\max_{ComAB} \frac{[(ComE \sim P)_D]}{[(ComE \sim P)_D] + K_{ComE \sim P_{AB}} * \left(1 + \frac{[ComE]^e}{Ki_{ComE_{AB}}^e}\right)} - \gamma_{ComAB} * [ComAB]$ |
| $\frac{d[ComD]}{dt} = \beta_{ComCDE} + v\max_{ComCDE} \frac{[(ComE \sim P)_D]}{[(ComE \sim P)_D] + K_{ComE \sim P} * \left(1 + \frac{[ComE]^e}{Ki_{ComE}^e}\right)} - 2 * k_{on\_D} * [ComD]^2 + 2 * k_{off\_D} * [(ComD)_D] - \gamma_{ComD} * [ComD]$ |
| $\frac{d[(ComD)_D]}{dt} = k_{on\_D} * [ComD]^2 - k_{off\_D} * [(ComD)_D] - v\max_{act} * \frac{[CSP]}{[CSP] + K_{act}} * [(ComD)_D] - \alpha_{auto} * [(ComD)_D]$ |
| $\frac{d[(ComD_{act})_D]}{dt} = v\max_{act} * \frac{[CSP]}{[CSP] + K_{act}} * [(ComD)_D] + \alpha_{auto} * [(ComD)_D] - \lambda * [ComE]^2 * [(ComD_{act})_D]$ |
| $\frac{d[(ComD_{consumed})_D]}{dt} = \lambda * [ComE]^2 * [(ComD_{act})_D] - \gamma_{ComD} * [(ComD_{consumed})_D]$ |
| $\frac{d[ComE]}{dt} = \beta_{ComCDE} + v\max_{ComCDE} \frac{[(ComE \sim P)_D]}{[(ComE \sim P)_D] + K_{ComE \sim P} * \left(1 + \frac{[ComE]^e}{Ki_{ComE}^e}\right)} + 2 * \rho * [(ComE \sim P)_D] - 2 * \lambda * [ComE]^2 * [(ComD_{act})_D] - \gamma_{ComE} * [ComE]$ |
| $\frac{d[(ComE \sim P)_D]}{dt} = \lambda * [ComE]^2 * [(ComD_{act})_D] - k_{on\_DprA\_EP} * [DprA]^2 * [(ComE \sim P)_D] - \rho * [(ComE \sim P)_D]$ |
| $\frac{d[ComX_{ina}]}{dt} = v\max_{ComX} * \frac{[(ComE \sim P)_D]^x}{[(ComE \sim P)_D]^x + K_{ComX}^x} + v\max_{DprA} * \frac{[ComX_{act}]^d}{[ComX_{act}]^d + K_{DprA}^d} + v\max_{SsbB} * \frac{[ComX_{act}]^s}{[ComX_{act}]^s + K_{SsbB}^s} - \omega_1 * [ComW] * [ComX_{ina}] - \gamma_{ComX} * [ComX_{ina}]$ |
| $\frac{d[ComX_{act}]}{dt} = \omega_1 * [ComW] * [ComX_{ina}] - v\max_{DprA} * \frac{[ComX_{act}]^d}{[ComX_{act}]^d + K_{DprA}^d} - v\max_{SsbB} * \frac{[ComX_{act}]^s}{[ComX_{act}]^s + K_{SsbB}^s} - \gamma_{ComX} * [ComX_{act}]$ |
| $\frac{d[ComW]}{dt} = v\max_{ComW} * \frac{[(ComE \sim P)_D]^w}{[(ComE \sim P)_D]^w + K_{ComW}^w} - \gamma_{ComW} * [ComW]$ |
| $\frac{d[DprA]}{dt} = v\max_{DprA} * \frac{[ComX_{act}]^d}{[ComX_{act}]^d + K_{DprA}^d} - 2 * k_{on\_DprA\_EP} * [DprA]^2 * [(ComE \sim P)_D] - \gamma_{DprA} * [DprA]$ |
| $\frac{d[(DprA\_ComE \sim P)_D]}{dt} = k_{on\_DprA\_EP} * [DprA]^2 * [(ComE \sim P)_D] - \gamma_{DprA} * [(DprA\_ComE \sim P)_D]$ |
| $\frac{d[SsbB]}{dt} = v\max_{SsbB} * \frac{[ComX_{act}]^s}{[ComX_{act}]^s + K_{SsbB}^s} - \gamma_{SsbB} * [SsbB]$ |
| $[ComD]_{total} = [ComD] + 2 * [(ComD)_D] + 2 * [(ComD_{act})_D] + 2 * [(ComD_{consumed})_D]$ |
| $[ComE]_{total} = [ComE] + 2 * [(ComE \sim P)_D] + 2 * [(DprA\_ComE \sim P)_D]$ |
| $[ComX]_{total} = [ComX_{ina}] + [ComX_{act}]$ |
| $[DprA]_{total} = [DprA] + 2 * [(DprA\_ComE \sim P)_D]$ |

The ODE approach provides detailed information on the network component dynamics, but requires high-quality data on component kinetics for estimating the parameters. However, in practice, quantitative measurements are often partial and available only for a fraction of the system’s entities. Therefore, some of the parameters of continuous models are usually based on inference. Here, the kinetic parameters of the regulatory network were assigned based on previous published experimental clues and on promoter activity measurements.

#### Estimations of protein synthesis rates and protein concentration kinetics

We exploited the data produced by Mirouze and collaborators, where transcriptional fusions of a luciferase reporter gene to the promoters of the target genes (*comC::luc*, *comX::luc*, *ssbB::luc*) were constructed in a wild type (WT) strain as well as in a *dprA*^−^ strain ((Mirouze *et al*, 2013) see Material and Methods therein). Real-time monitoring of gene expression was conducted when competence was induced by addition of a saturating concentration of exogenous CSP that allows synchronization of the population response. Luminescence (expressed in relative luminescence units, RLU) and absorbance (OD_492nm_) values were recorded every minute after CSP addition over a period of 45 minutes. The quantity of luminescence per cell as a function of time (*r*(*t*)) was calculated as the ratio *r*(*t*)) = *I*(*t*)/*A*(*t*), where *I*(*t*) is the luminescence intensity (in RLU) and *A*(*t*) the absorbance values corrected by subtraction of the absorbance background measured on wells containing only growth medium (de Jong *et al*, 2010). Since the luciferase does not require any post-translational modification such as folding, this ratio estimates the average concentration of reporter protein per cell. Thus, the dynamics of the system is conveniently described by the temporal evolution of the luciferase concentration.

The next step was to transform the normalized luminescence signal per cell into promoter activities and protein kinetics by following the mathematical model developed by Stefan and collaborators (Stefan *et al*, 2015). To calculate the evolution of the concentration of a given protein over time, a correction was performed to take into account the differences in half-lives between the reporter luciferase and the protein whose gene activity is measured. For ComX and SsbB, which have a shorter life times (estimated at 8 min, *t_1/2_* = 5.45 min) than luciferase (measured at 21.6 min, *t_1/2_* =15 min), the uncorrected values clearly overestimate the protein concentration (Appendix Figure S2). Conversely, for ComE and ComD whose life-time has been assessed to be 80 min (*t_1/2_* = 55.45 min), the uncorrected values underestimate the protein concentration (Appendix Figure S2).

The kinetics of promoter activities and protein concentration kinetics deduced from the luminescence data of *comC::luc*, *comX::luc* and *ssbB::luc* in both wild type and *dprA*^−^ cells are shown in Figure 2. For wild type cells, a promoter activity pulse for each studied promoter is observed following CSP addition in the medium, consistent with the fact that competence is a transient physiological state. The curve of the calculated protein kinetics presents a shape similar to that of the promoter activity curve, with a shift owing to the time required to translate the mRNAs into proteins and a decreasing slope that is consistent with the protein half-lives, *i. e.*, steeper for ComX and SsbB with short half-lives than for ComD and ComE. In *dprA*^−^ cells, coherence between the deduced protein synthesis rates and the different promoter activities is also observed. The alteration of *comX* transcription shut-off in the *dprA* mutant is clearly observed as is its incidence on the maintenance of *ssbB* expression. Moreover, our estimates of protein concentration kinetics obtained in the wild type strain are in agreement with published experimental results, supporting our choice of protein half-life values for which no precise measurements are available. For ComD and ComE, the concentration peak is reached about 15 min after CSP addition and the level remains stable over a period of about 25 min, in accordance with the Western blot results obtained by Martin and collaborators (Martin *et al*, 2013). ComX appears between five and ten min after CSP addition, reaches a concentration peak after 15 min and then decays. These kinetics fit the published experimental data (Piotrowski *et al*, 2009; Luo & Morrison, 2003). SsbB kinetics are also consistent with previously published results (Mirouze *et al*, 2013), with the detection of the protein about 5 min after competence induction, a maximum concentration around 15-20 min and an almost complete disappearance of protein after 60 min. Finally, the prolongation of ComX presence in a *dprA*-context as shown by Weng and collaborators (Weng *et al*, 2013) is reproduced, with the same temporality.

**Figure 2.**
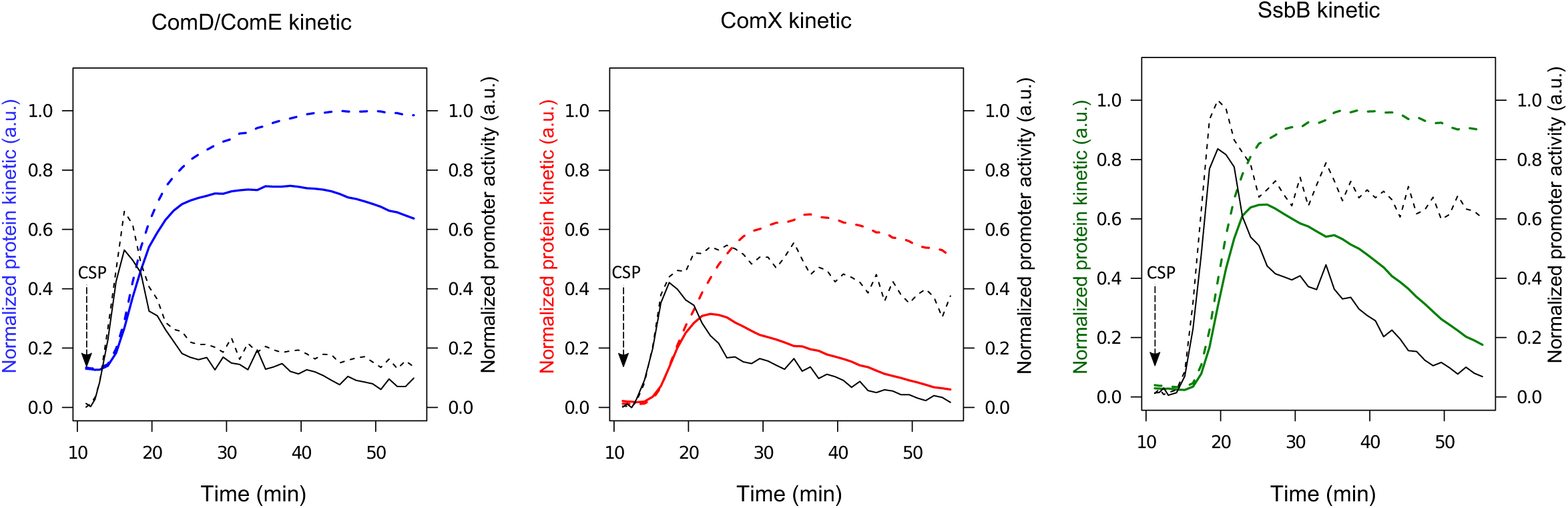
Promoter activities and protein kinetics deduced from luminescence data. Promoter activities per cell computed from normalized raw luminescence signals are represented by black lines (right scale). Reconstructed protein kinetics corrected for differences in half-lives between luciferase and the protein whose gene activity is measured are represented by colored lines (left scale). Solid and dashed lines correspond to computed values obtained from raw luminescence data in WT and *dprA*^−^ strains respectively (Mirouze *et al*, 2013). The protein life times used for computation are: 8 min for ComX and SsbB, 80 min for ComD and ComE and 21.6 min for luciferase(Prudhomme & Claverys, 2007). The *comD* and *comE* genes being in an operon, the blue line represents both ComD_total_ as ComE_total_ kinetics. Red and green lines depict ComX_total_ and SsbB kinetics respectively. Promoter activities and protein concentrations have been normalized with respect to the maximum values obtained over all computed data sets; therefore values are given in arbitrary units (a.u.).

#### Parameter estimation

The parameter values were estimated by exploiting published data where they were available. Using results from gel-shift experiments indicating that the phosphorylmimetic ComE^D58E^ has a greater affinity than ComE for P_*comC*_ and that P_*comC*_ is a stronger promoter than P_*comX*_ (Martin *et al*, 2013), we set up constraints on the parameter values with *K_comX_*>*K_comE_*>*K_comE∼p_*. As no accurate protein life time measurements were available, intervals were determined for degradation rate constants based on Western blot analyses: [4,12] min^−1^ for unstable proteins ComX and SsbB and [40,120] min^−1^ for stable proteins ComE, ComD and DprA (Martin *et al*, 2013; Mirouze *et al*, 2013; Weng *et al*, 2013). We also exploited the experimental results obtained from the detailed studies on the AgrCA TCS of *S.aureus.* As ComD possesses 50% similarity with AgrC and ComE 52% with AgrA, we can assume that both TCSs will share similar biochemical properties. A half-life of 3.9 min has been measured for the phosphorylated form of the response regulator AgrA, corresponding to an average life time of 5.26 min and to a self-catalyzed dephosphorylation rate of 0.18 min^−1^ (Wang *et al*, 2014). Therefore, the search space for estimating ComE∼P dephosphorylation rate *ρ* has been restricted to the interval [0.1,0.4] min^−1^. The half-life of the phosphorylated form of AgrC (66 seconds) reveals an efficient phosphoryl group transfer between AgrC∼P and AgrA corresponding to a transphosphorylation rate of 0.63 min^−1^(Wang *et al*, 2014). Thus, the search space for estimating transphosphorylation constant *λ* was restrained to [0.4,1] min^−1^.

Initial protein concentrations were chosen such that the system is at steady-state at the beginning of the simulation, corresponding to the non-competence/vegetative state, and such that the addition of exogenous CSP will initiate competence development.

Three datasets were used concomitantly to estimate the parameter values: the two previously described time-series luminescence signals obtained from WT and *dprA*^−^ strains (Mirouze *et al*, 2013) transformed into protein kinetics as previously described, and a third corresponding to the protein kinetics in a *clpP*^−^ strain. In the latter case, no luminescence data were available but cells have been shown to escape from the competent state as rapidly in a *clpP* mutant as in a wild-type strain (Piotrowski *et al*, 2009). Thus, we generated the *clpP* mutant kinetic data, where ComX is stable, using the WT dataset and setting the degradation rate constant *γ_ComX_* of ComX to 0 for inferring ComX kinetics.

Parameters were estimated using the particle swarm optimization (PSO) method (Eberhart & Kennedy, 1995), as it had been shown to perform the best (Baker *et al*, 2010). The objective function that was minimized corresponds to the mean square distance between the calculated experimental protein concentration kinetics (ComD, ComE, ComX and SsbB) and model estimated protein concentration kinetics ([ComD]_total_, [ComE]_total_, [ComX]_total_ and [SsbB]). To generate protein concentration kinetics for the *dprA* mutant, the maximal transcription rate of *dprA* (*vmax_dprA_*) was set to 0, while for the *clpP* mutant both degradation constants of ComX (*γ_comX_*) and ComW (*γ_comW_*) were set to 0.

#### Validation of the ODE model

Comparison of simulated data with the experimental measurements made on the three strains (WT, *dprA*^−^ and *clpP*^−^) reveals a relatively good agreement (Figure 3). However, discrepancies between the measured and simulated values are observed, especially in the case of SsbB kinetics. Moreover, in the *clpP* mutant, the steady decrease in SsbB seen 20 min after CSP addition, a marker of competence shut-off, is not reproduced by our simulation, where the level of SsbB remains stable over more than 50 min. Therefore, we conclude that another, as yet unknown, actor is involved in late gene transcription termination, as proposed by Piotrowski and collaborators (Piotrowski *et al*, 2009) and Weng and collaborators (Weng *et al*, 2013).

**Figure 3.**
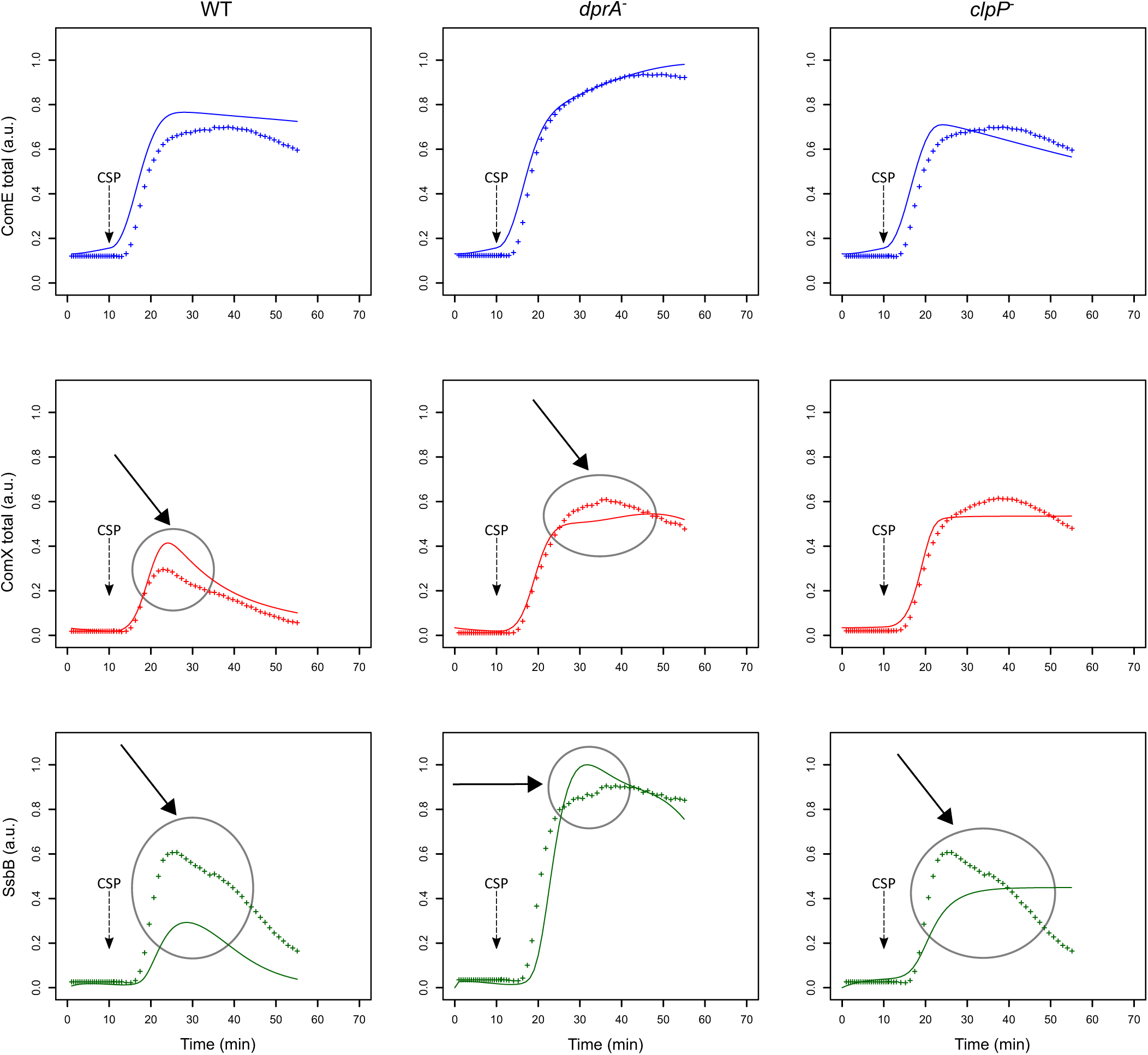
Comparison of experimental and simulated protein kinetics obtained with the initial model. Comparison of simulated data with the experimental measurements are shown for the WT strain, the *dprA* mutant strain and the *clpP* mutant strain. Since luminescence data was not available for the *clpP* mutant, they were generated using the WT dataset and ComX kinetics was inferred by setting ComX degradation constant *γ_comx_* to 0. For the two other strains, reconstructed protein kinetics correspond to those of figure 2. Experimental and simulated data are represented by crosses and solid lines respectively. In the *dprA* mutant the simulated protein kinetics were obtained by setting the DprA maximal synthesis rate (*vmax_dprA_*) to 0. In the *clpP* mutant both ComX (*γ_comx_*) and ComW (*γ_comw_*) degradation constants were designated as 0. Competence gene expression was assumed to be at steady-state at the beginning of the simulation, and competence development was induced by adding one arbitrary unit (a.u.) of CSP (corresponding to 100 ng/mL) at *t*= 10 min in order to reproduce the experimental protocol (100 ng/mL added after 10 min incubation (Mirouze *et al*, 2013)). The same color code as in Figure 2 is used. Major discrepancies between experimental and simulated data are circled and indicated by an arrow.

#### Optimization of the initial ODE model

Consequently, we have modified this initial model to integrate a new unknown gene, which we name *comZ*, whose synthesis could be under the control of either ComE∼P (early *com* gene) or ComX (late *com* gene). We considered four alternative hypotheses for ComZ action (Appendix Figure S3): i) ComZ interacts with ComW and thus prevents its action in the transition from the inactive to the active form of ComX, ii) ComZ competes with ComW for interaction with the inactive form of ComX and affects the formation of the ComX active form, iii) ComZ inhibits the active form of ComX directly, and iv) ComZ and the active form of ComX compete for binding to RNA polymerase, in which case ComZ would correspond to the *σ*^A^ factor. For each of the eight models, the changes generated in the ODEs are reported in Appendix Table S1. The parameter estimations and the network simulations were performed as previously. Since the models cannot be compared directly on the basis of their objective function value, we used the Akaike Information Criterion (AIC). AIC was computed for each candidate model (Table 3) and the one having the smallest AIC value was selected since it is considered to be the closest to the unknown reality that produces the data. This corresponds to the model in which ComZ is a late-gene product that interacts directly with ComW, impairing the formation of the active form of ComX. Validation of this model is described in more detail below and is further tested to check its predictive behavior in the specific experimental conditions used. Comparison of simulated data with the experimental measurements made on the three strains (WT, *dprA*^−^ and *clpP*^−^) for the other models are shown in Appendix Figures S4 to S10. Clearly, direct inhibition of the active form of ComX by ComZ, whether an early or late *com* gene product, can be ruled out since the simulated values reproduce poorly the experimental kinetics, especially for SsbB (Appendix Figures S9 and S10). Moreover, for the same hypothetical action of ComZ, the lowest AIC values are obtained with models based on the hypothesis that ComZ is a late *com* gene product, and therefore better reproduce the experimental data (Table 3).

**Table 3.** Akaike’s Information Criterion (AIC) computed for each candidate model

| Alternative models | RSS<sup>&</sup> | Parameter | AIC<sup>&&</sup> |
| --- | --- | --- | --- |
| Interaction between ComW and a late *com* gene product ComZ impairs ComW activity (Figure 4) | 0.33 | 44 | -5760.95 |
| Competition between the active form of ComX and a late *com* gene product ComZ for RNA polymerase binding (Figure S4) | 0.44 | 50 | -5522.92 |
| Competition between ComW and a late *com* gene product ComZ for the inactive form of ComX impairs the formation of the active form of ComX (Figure S5) | 0.54 | 47 | -5382.64 |
| Interaction between ComW and an early *com* gene product ComZ impairs ComW activity (Figure S6) | 0.63 | 44 | -5272.10 |
| Competition between the active form of ComX and an early *com* gene product ComZ for RNA polymerase binding (Figure S7) | 0.74 | 48 | -5137.34 |
| Competition between ComW and an early *com* gene product ComZ for the inactive form of ComX impairs the formation of the active form of ComX (Figure S8) | 1.58 | 47 | -4572.90 |
| Inhibition of the active form of ComX by a late *com* gene product ComZ (Figure S9) | 2.47 | 44 | -4239.21 |
| **Initial model based on current biological knowledge without involving an additional ComZ partner (Figure 3)** | **2.63** | **39** | **-4220.38** |
| Inhibition of the active form of ComX by an early *com* gene product ComZ (Figure S10) | 3.32 | 44 | -4015.63 |

<sup>&</sup> Residual squared sum<sup>&&</sup> $AIC = n \ln\left(\frac{RSS}{n}\right) + 2k$ where $k$ is the number of estimated model parameters, $n$ is the number of observations (756) and $RSS$ is the sum of squared residuals of the fitted model

#### Validation of the new selected model: an unknown late gene product ComZ interacts with ComW and impairs its action

This new model gives good agreement between the simulated and measured kinetics (Figure 4), not only for the wild type strain but also for the *dprA* and *clpP* mutants. Indeed, the value of 0.33 obtained for the objective function value, measuring the sum of squared errors between measurements and model predictions, is very small. Moreover, despite the stabilization of ComX in the *clpP* mutant, the kinetics of SsbB reveal that this cell escapes the competent state as rapidly as the wild type cell.

**Figure 4.**
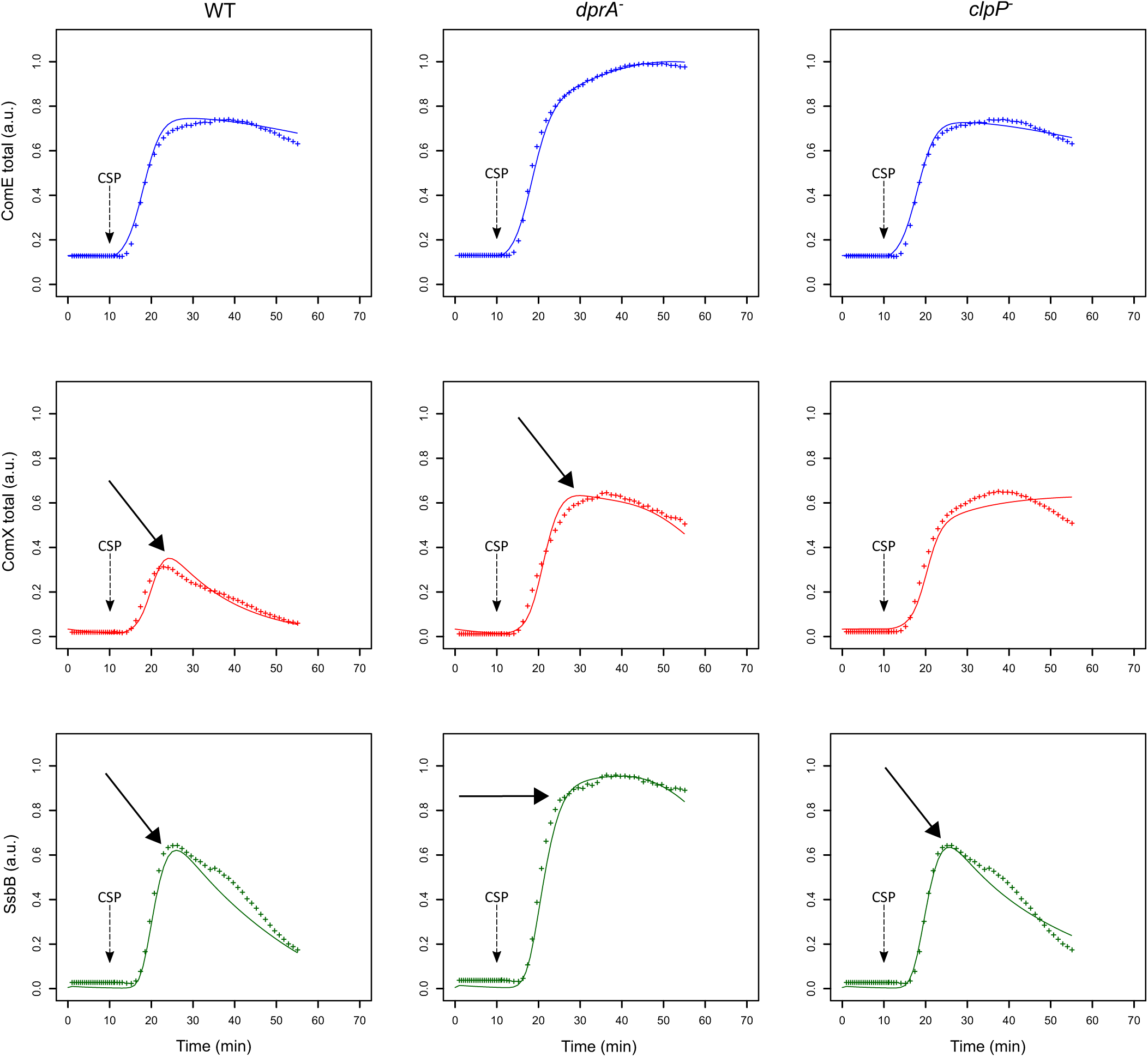
Comparison of experimental and simulated protein kinetics obtained with the modified selected model. An unknown late gene product ComZ interacting with ComW and preventing the transition from the inactive to the active form of ComX has been introduced in the initial model. Experimental and simulated conditions are the same as in Figure 3 as is the color code. Arrows indicate the major discrepancies between experimental and simulated data observed in Figure 3, showing that the new model greatly enhances the fitting between both curves.

The estimated parameter values (Table 4) appear in accordance with experimental measurements. Our estimated degradation rate constants lead to a half-life of ComE, ComD and DprA of 83 min, which have been shown experimentally to be stable over a period of at least 80 min (Martin *et al*, 2013; Mirouze *et al*, 2013; Weng *et al*, 2013). The estimated halflives of ComX and comW at 8 min, and that of SsbB at 6 min are close to their experimental values of around 5 min (Mirouze *et al*, 2013; Piotrowski *et al*, 2009). The estimated value of K_ComE∼p_ (0.15), the binding constant for ComE∼P to P_*comC*_, shows about a threefold reduction compared to the estimated value of Ki_ComE_ (0.44), the binding constant for ComE to P_*comC*_. This is about the same order of magnitude as the fourfold reduction of the apparent Kd for *P_comC_* reported for the phosphorylmimetic mutant compared to the Kd of ComE (Martin *et al*, 2013). The estimated binding constant (K_comX_) for ComE∼P to P_*comX*_ shows an increase of 29-fold compared to that for P_*comC*_. This is slightly higher than the average tenfold increase reported by Martin and collaborators (Martin *et al*, 2013). Finally, our estimated half-life of ComE∼P is 2.23 min, comparable to 3.9 min measured for the phosphorylated form of AgrA (Wang *et al*, 2014), while the value of the transphorylation rate constant λ has been estimated at 1 min^−1^ compared with 0.63 min^−1^ for the phosphoryl group transfer constant in the TCS AgrAC (Wang *et al*, 2014).

**Table 4.**
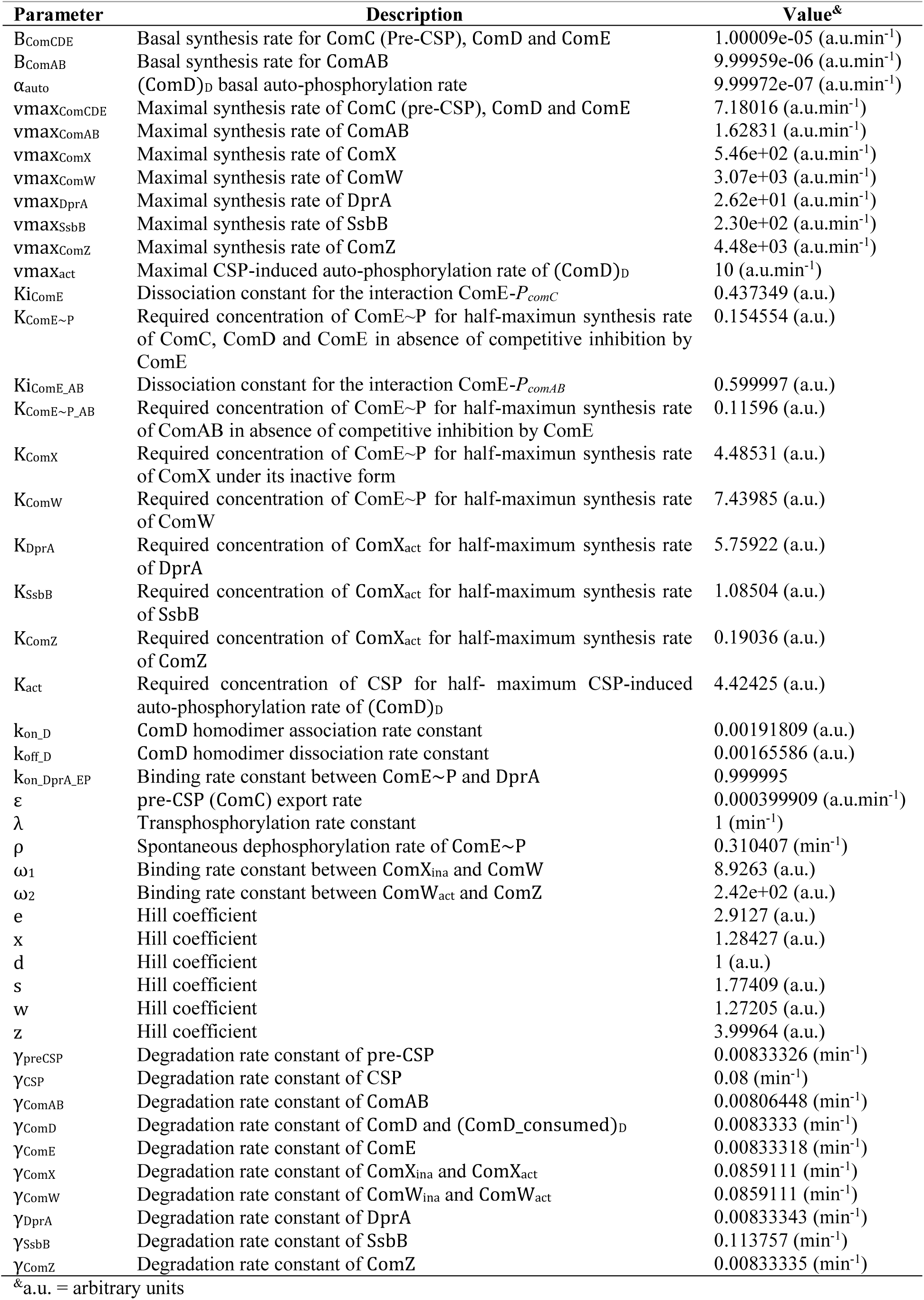
Description and numerical values of parameters from the selected alternative model

#### Both CSP-ComD interaction and ComD autophosphorylation are pH sensitive

To further challenge our model, we tested the impact of the initial pH of the growth medium on competence development, since it was reported that this is an important parameter in controlling competence (Chen & Morrison, 1987). However, this effect has never been studied experimentally in detail. In the total absence of CSP, basal expression of the master competence operon *comCDE* depends on an intact ComD-ComE TCS, meaning that ComD basal auto-phosphorylation and phosphate transfer to ComE occurs (Martin *et al*, 2010). The initial pH of the growth medium may affect either ComD basal auto-phosphorylation or the kinetics of autophosphorylation of ComD bound to CSP (CSP-induced phosphorylation). To distinguish between these options, we recorded *comCDE* promoter activity during the first chain of induction from CSP to ComE∼P in a strain that does not allow export or expression of CSP (see Material and Methods). The cells were first grown to allow a basal expression of ComDE to reach the same non-competent physiological state. Samples were then transferred to growth medium of different initial pH (ranging from 6.8 to 8.19) and different CSP concentrations (0, 25, 50 and 100 ng.ml^−1^). RLU was directly recorded using the luciferase gene under control of either the *comCDE* promoter (strain R1205) or the tRNA^arg^ gene promoter located just upstream of the *comCDE* operon but not part of the competence regulon (strain R1694) (Martin *et al*, 2010).

In the absence of CSP, the increase of the luminescence values over a period of 20 min appears linear for both strains (competence and non-competence reporters) and was independent of initial pH of the growth medium. The slope of the curve was computed for each pH value from the average of the luminescence values measured on four replicates at each time point. A slight increase of the slope is observed depending on the pH value. This increase appears similar for the competence and non-competence reporter strains (Figure 5A), and much lower than that obtained when the initial slope was computed for the experiment in which 36 ng.ml^−1^ of CSP is added (Figure 5A). Since no convincing difference was observed between the results obtained without CSP addition on the strain R1205 and the R1694 control strain, the hypothesis that ComD alone could be a sensor of pH can be eliminated.

**Figure 5.**
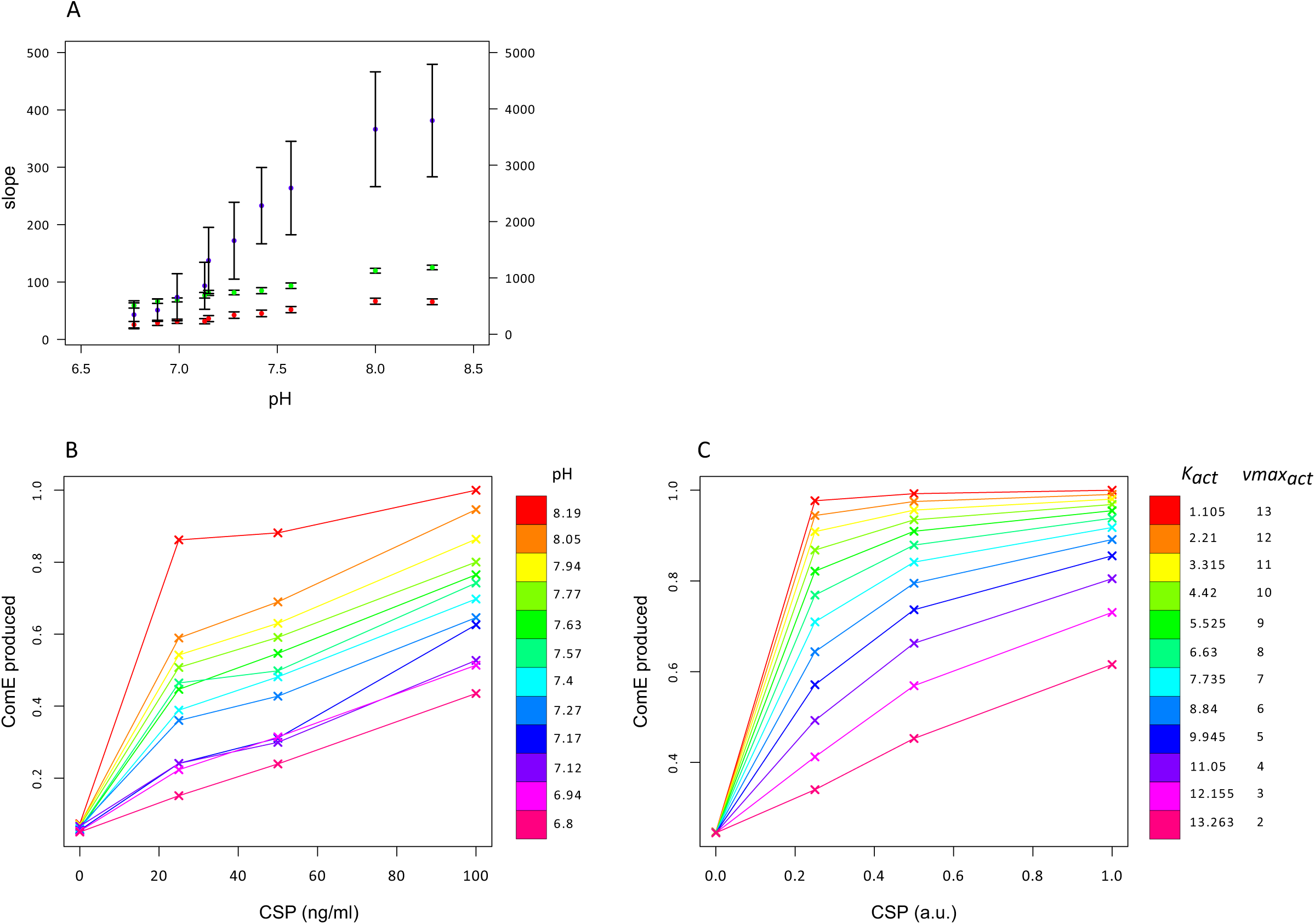
The initial pH value of the growth medium affects the comCDE circuit. (A) For a given pH value of the growth medium, each red and green point corresponds to the slope of the curve computed from the average of the luminescence values measured on the four replicates at each time point over the first 20 min of the experiment for the strain R1205 (red) and the R1694 control strain (green) respectively (left scale). The purple points correspond to the initial slope of the curve calculated over the first 3 min of the experiment for each pH value when 36ng/ml of CSP is added in the medium (right scale). The confidence interval on the slope is plotted. (B) Each point of the curves corresponds to the quantity of ComE synthesized over a period of 20 min after CSP addition for each CSP-pH combination tested experimentally. This quantity is obtained by calculating the area under the curve of the kinetics of ComE and is normalized with respect to the maximum value of ComE synthesized over all experiments. CSP concentrations used in the experiments: 0, 25, 50 and 100 ng/mL. Strain used: *P_comc_::luc* R1205. (C) Each point of the curves corresponds to the simulated quantity of ComEtotai synthesized over a period of 20 min after CSP addition. Protein amounts are calculated as in (B) and normalized with respect to the maximum value of ComE synthesized over all simulations. Simulations are performed by tuning the two ODE parameters corresponding to the affinity of a ComD dimer for the CSP (*K_act_*) and CSP-induced autophosphorylation rate of ComD dimer (*vmax_act_*). CSP amounts used in simulations: 0, 0.25, 0.5, 1 a.u. *comA*^−^ is simulated by setting *β_comAB_*= 0 (basal synthesis rate of ComAB) and *vmax_comAB_* = 0 (maximal synthesis rate of ComAB).

The luminescence data were processed for each CSP-pH combination, as previously, to obtain the average kinetics of ComE concentration per cell. The amount of ComE produced over a period of 20 minutes immediately after CSP addition was then obtained by calculating the area under the curve of the ComE time courses. We assumed that this value is proportional to the initial rate of ComE accumulation and therefore to ComE synthesis rate. The results obtained when CSP is used to induce the system (Figure 5B) clearly show that the initial pH of the growth medium acts on the efficiency of competence development. For acidic pH values (≤ 7.12) and at a CSP concentration of 25 ng.ml^−1^ very small quantities of ComE are synthesized, and even at the highest CSP concentration (100 ng.ml^−1^) the level of ComE reaches only about half that obtained at pH 8.19. In particular, a higher quantity of ComE is obtained for cells induced with 25 ng.ml^−1^ of CSP at pH 7.57 than for cells induced with 100 ng.ml^−1^ of CSP at pH 6.8. For a given CSP concentration, the more alkaline the medium, the higher the amount of ComE synthesized. This effect is more pronounced at pH 8.19 where ComE rapidly attains a plateau at a CSP concentration of 25 ng.ml^−1^. Those results clearly show that the pH of the growth medium is an important parameter in controlling competence development.

To determine whether the model can simulate these experimental data, we assumed that the pH of the growth medium affects reactions taking place in the extra-cellular environment, namely: i) interaction between the CSP and ComD and its effect on ComD phosphorylation (Table 1, reaction (5)), or ii) the CSP degradation rate (Table 1, reaction (12). Since, HtrA protease activity, involved in CSP degradation, appears unaffected by pH (Cassone *et al*, 2012), we deduced that the CSP decay rate is not sensitive to pH variation. Therefore, to mimic the pH variation, we performed a parameter sweep, jointly, on the affinity of a ComD dimer for CSP (*K_act_*) and the maximum rate of CSP-stimulated conversion of the unphosphorylated form of a ComD dimer into its phosphorylated form (*vmax_act_*). Indeed, tuning only the values of one of these two parameters does not reproduce the experimental data. To establish the parameter scale, we set the values of *K*_*ac*t_ and *vmax_act_* at pH 7.8 to those previously estimated (4.42 and 10 respectively) for wild-type and *dprA*^−^ cells since this pH value was used in these experiments (Mirouze *et al*, 2013). This parameter sweep was achieved for each of the four CSP concentrations used to induce the system. Each point on the curves corresponds to the simulated quantity of ComE synthesized over a period of 20 min after CSP addition, obtained as before by calculating the area under the curve of the ComE time courses.

We observe good qualitative agreement between experimental and simulated curves (Figure 5C). In particular, the shape of the experimental curve obtained for the most acidic pH value (6.8) was reproduced with an amount of ComE that increases linearly and reaches a maximum of about half the level obtained with the *K_act_* and *vmax_act_* values corresponding to growth medium with the highest alkaline pH. In the latter case, the simulated values of ComE rapidly attain a plateau at low CSP concentrations as observed experimentally at pH 8.19. Thus, by adapting the values of two ODE model parameters involved in phosphorylation of ComD through its interaction with CSP, we could reproduce the observed experimental data. This result further validates our modeling and predicts that the initial pH of the growth medium affects the reaction controlling the activation of ComD through CSP-induced phosphorylation.

#### Investigating the model’s predictive behavior on new aspects of the competence cycle

We next explored how the model predicted the effect of the basal rates of early competence gene expression on the spontaneous shift to competence and how it can reproduce and explain the blind-to-CSP period.

#### Coordinated rates of ComCDE and ComAB basal expression are required to elicit spontaneous competence

Although the model was developed in a context of competence induction by external CSP, we investigated whether it could simulate the triggering of spontaneous competence. We first tested the basic idea that over-expression of *comCDE* should lead to derepression of competence. We followed the network dynamics without adding external CSP for system induction and we performed a parameter sweep over the ComCDE basal synthesis rate (from 10^−5^ to 10^−2^ a.u. min^−1^). Contrary to expectation, this basal rate increase does not activate competence but inhibits competence initiation (Appendix Figure S11A). Computation of protein and peptide amounts at *t* = 20 min clearly shows that ComE and intracellular pre-CSP increase according to the rise in basal synthesis rate whereas the level of mature exported CSP remains constant and very low (from 4.3e-^05^ to 5.9e-^05^ a.u.), such that ComX and SsbB are not synthesized (Figure 6). However, when we increase the ComCDE and ComAB basal synthesis rates simultaneously, competence is restored as shown by appearance of SsbB production at *t* = 20 min (Figure 6 and Appendix Figure S11B for protein and peptide level kinetics). We deduce from this result that the export of CSP through ComAB is limiting, as observed experimentally (Martin *et al*, 2000), leading to an excess of ComE over ComE∼P that prevents ComE∼P from competing with ComE for binding to P_*comC*_.

**Figure 6.**
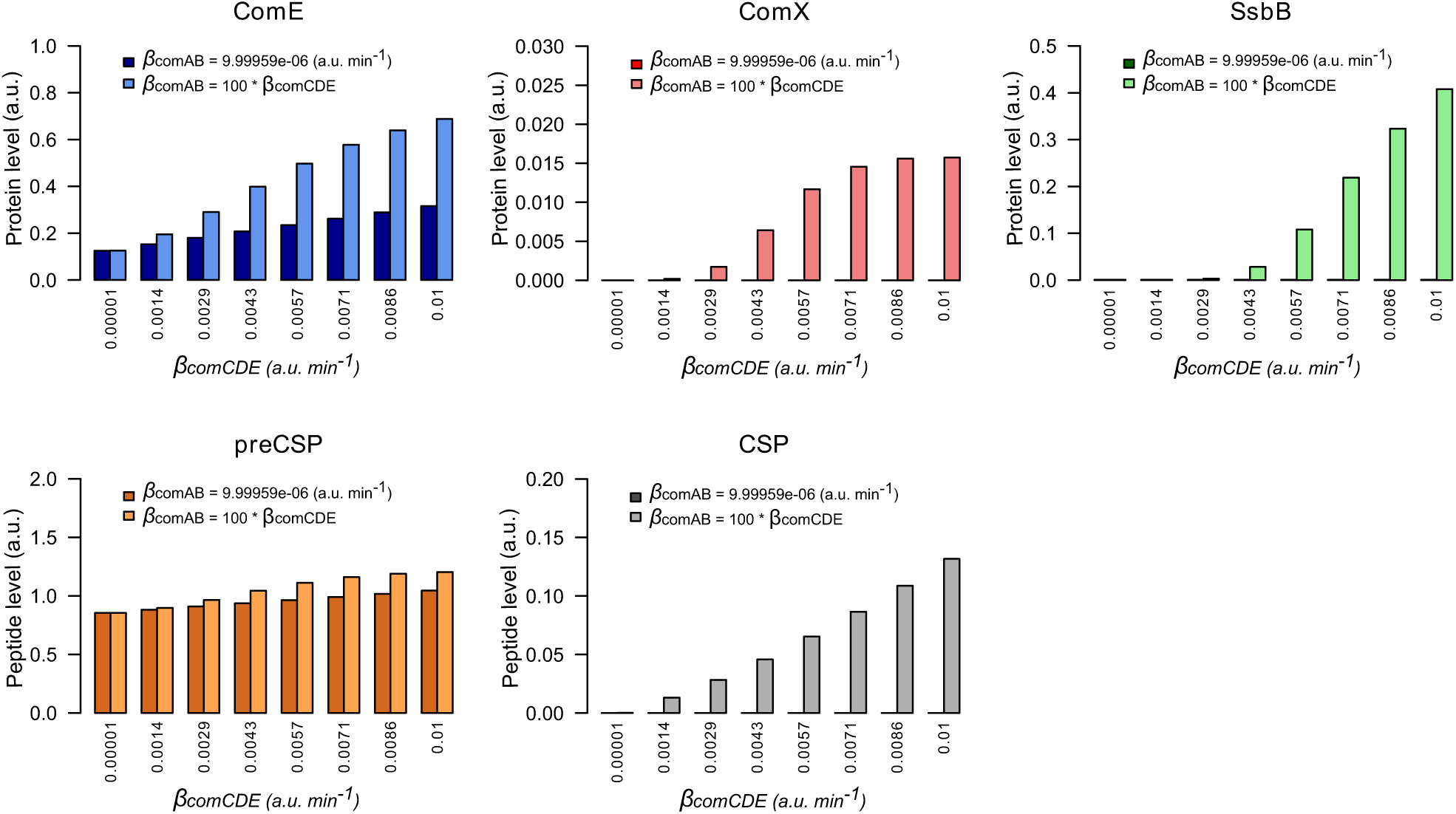
Both basal synthesis rates of *comAB* and *comCDE* influence spontaneous competence development. Comparison of the protein and peptide levels computed à *t* = 20 min when the simulated network dynamics is followed without external CSP addition for a ComCDE basal synthesis rate (*β_comCDE_*) varying from 10^−5^ to 10^−2^ a.u. min^−1^. Dark color bars show the results obtained when the exporter basal synthesis rate (*β_comAB_*) keeps a constant value corresponding to that estimated in our model (Table 4). Light color bars display the values obtained when *β_comAB_* varies conjointly with *β_comCDE_* with *β_comAB_* = 100 ^∗^ *β_comCDE_*.

Then, for a given value of the basal synthesis rate of ComCDE (0.005 a.u. min^−1^), we gradually increase the ComAB basal synthesis rate from 0.01 min^−1^ to 1 min^−1^. Significant synthesis of SsbB was observed when the basal ComAB synthesis rate reached 0.05, indicating that activation of the competence state can be restored (Appendix Figure S12). However, as the basal ComAB synthesis rate decreases, we observe an increasing delay in competence triggering as estimated by the time required for SsbB to exceed 0.1 a.u. (Figure 7). For example, this threshold is exceeded at 18 min with a basal ComAB synthesis rate of 1 and only at 38 min with a basal synthesis rate of 0.05. This is consistent with a more efficient processing of pre-CSP that allows extracellular CSP to reach the critical concentration required to activate ComDE, and hence more rapid setting up of the positive feedback loop (Appendix Figure S12). The ratio ComE∼P/ComE at time *t* when SsbB exceeds 0.1 a.u., computed for each simulation, varies from around 4e^−03^ to 6e^−03^. We propose that below this value, the level of ComE∼P is insufficient to set up the positive feedback loop. All these simulations suggest that a balance between the basic synthesis rates of ComCDE and ComAB is crucial to triggering spontaneous competence.

**Figure 7.**
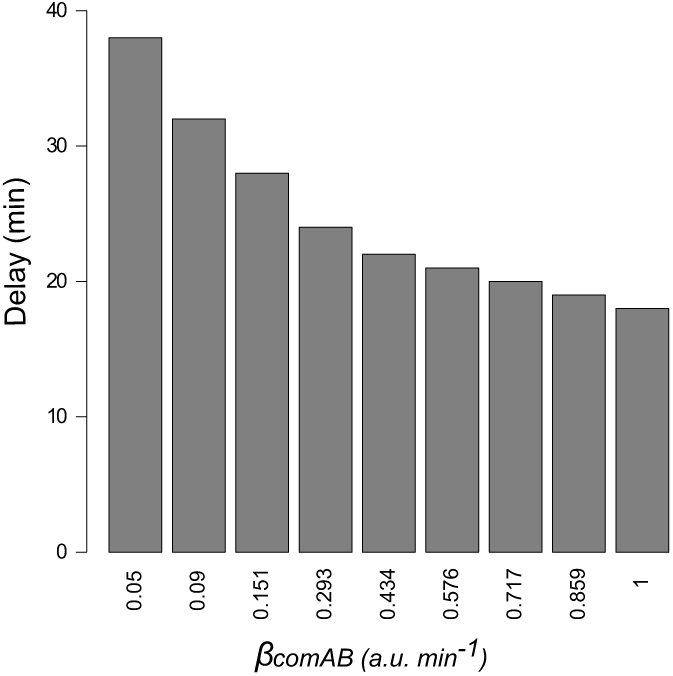
Efficient export of matured pre-CSP promotes establishment of the positive feedback loop. Time delay measured for competence triggering estimated as the time required for SsbB to exceed 0.1 a.u. in network simulations where, for a fixed value of the ComCDE basal synthesis rate (0.005 a.u. min^−1^), the ComAB basal synthesis rate (*β_comAB_*) is gradually increased from 0.01 min^−1^ to 1 min^−1^. For *β_comAB_* equal to 0.01 and 0.02 min^−1^, SsbB does not exceed the defined threshold.

#### Blind-to-CSP period results from the blocking of the ComD/E pathway through DprA accumulation

After the shut-off phase, the cells are unresponsive to higher concentrations of CSP for 60-80 min, the refractory- or blind-to-CSP period (Chen & Morrison, 1987; Hotchkiss, 1954). It was suggested that the stability of DprA could explain this insensitivity (Mirouze *et al*, 2013). For testing whether the model can reproduce the blind-to-CSP period, we simulated this situation by deducing the consequences of adding CSP to the medium at different times after competence shut-off estimated at *t* = 50 min (Figure 8A). We did not obtain an on/off response, but over a period of 80 min after competence shut-off the response of the cell to CSP addition is less efficient. We explain this result by the presence, after competence shut-off, of free residual DprA (not involved in a complex with ComE∼P) and an excess of ComE. After a second CSP addition, the residual ComE and newly synthesized ComE are phosphorylated by a ComD dimer. However, in the model, the level of residual DprA is insufficient to sequester all the available ComE∼P and a second wave of competence can be initiated. The magnitude of this wave increases progressively with time elapsed since the previous competence shut-off, and by 120 minutes equals that of the first wave. This is consistent with the estimated 120 min effective stability of DprA in the model (half-life of 83 min). To corroborate this hypothesis, we performed two additional simulations. In the first, we removed all residual DprA before adding CSP at the shut-off phase. This precludes the refractory period and a second peak of competence identical to the first is observed (Figure 8B). In the second simulation, we increased the DprA level in the cell before the second addition of CSP by changing its current value of 1.35 a.u. into 2.7 a.u. It this case, the cell is totally unresponsive to the CSP (Figure 8C). Therefore, our simulated data strengthen the hypothesis that DprA is a main actor in the blind-to-CSP period.

**Figure 8.**
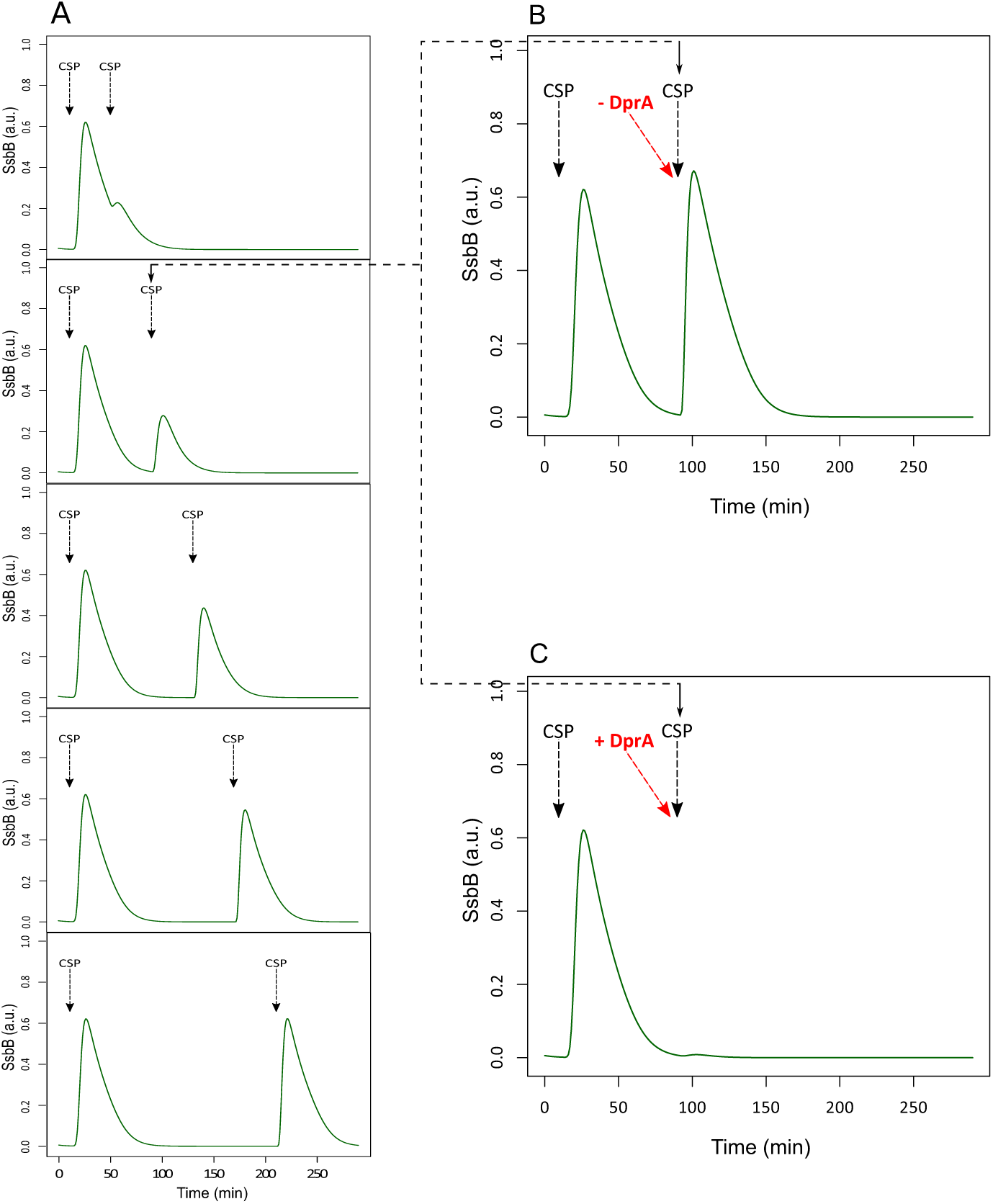
DprA has a major role in the blind-to-CSP period. (A) SsbB kinetics obtained in five independent simulations. In each simulation, the external CSP is added twice: at time *t* = 10 min in all experiments and at time *t* = 50, 90, 130, 170, 210 in the experiments corresponding respectively to 0, 40, 80, 120 and 160 min after competence shut-off. (B) Simulated SsbB kinetics where competence is initially induced by addition of external CSP at 10 min and the system cleared of residual DprA protein one minute before the second addition of CSP at *t* = 90 min, *i.e.*, 30 min after competence shut-off. The removal of residual DprAs is achieved by changing its current protein level (1.35 a.u.) into its estimated initial value in our model (0.25 a.u.). (C) Simulated SsbB kinetics obtained as in (B) but where the level of DprA is increased one minute before the second addition of external CSP by changing its current value (1.35 a.u.) to 2.7 a.u. The dotted line indicates the second addition of CSP at *t* = 90 min.

To confirm that the refractory period results only from regulation of the early *com* genes by the ComE/ComE∼P ratio, we modified our model to bypass the activation of *comX* and *comW* by ComE∼P. In the ODEs describing the kinetics of ComX and ComW we included a constant synthesis rate that allows a direct induction of the gene. In such a model, ComX and ComW can be produced either by the ComE∼P pathway or independently by direct induction of both genes. We ran simulations by adding either CSP or by directly activating *comX* and *comW* at the end of the shut-off phase (Figure 9A). While the model is almost unresponsive to the second addition of CSP, it can develop a second wave of competence after direct activation of *comX/comW* despite the presence of the late protein DprA (Figure 9A). To experimentally validate our prediction, we took advantage of the previously described inducible CEPr expression platform (supplementary information in (Johnston *et al*, 2016)). This platform is controlled by the BlpR/H two-component system (de Saizieu *et al*, 2000) which is activated by the mature bacteriocin induction peptide (BIP). *comX* and *comW* genes were cloned as an operon under control of BlpR and introduced into strain R3584 carrying a luciferase gene expressed from the *ssbB* promoter. This strain, R3932, can produce ComX and ComW either by the ComE∼P pathway or independently by the BlpR/H system. During the competence shutoff, only BIP addition allows new expression of late competence genes (Figure 9B) as predicted by the model. These results strongly support the conclusion that the blind-to-CSP period results from blocking of the ComD/E pathway by DprA accumulation which prevents reappearance of a ComE∼P free pool.

**Figure 9.**
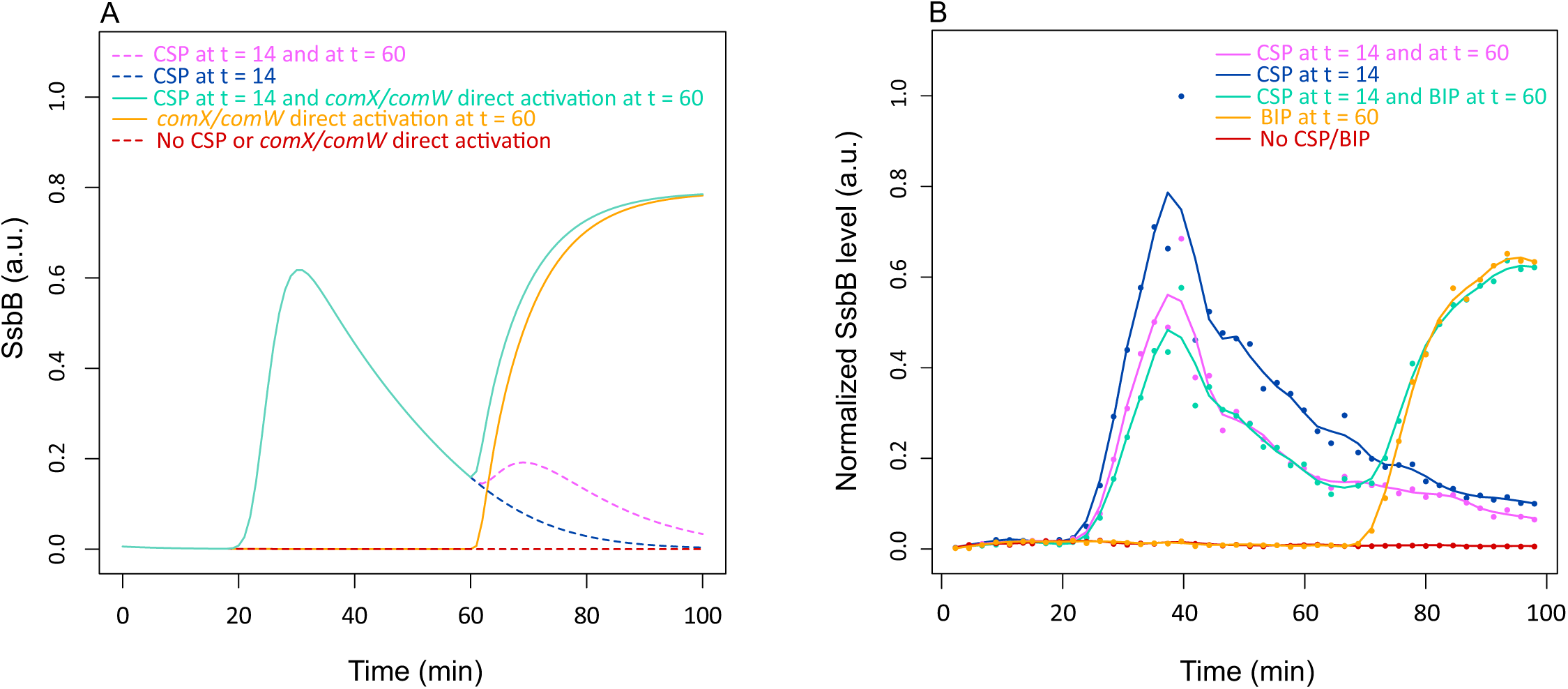
Bypassing the ComD/E pathway for ComX and ComW synthesis suppresses the blind-to-CSP period. (A) Simulation of SsbB kinetics by the modified model, allowing the synthesis of ComX and ComW either dependent on or independent of the ComCDE regulatory pathway. The simulation of the direct induction of ComX and ComW synthesis is achieved by shifting the value of the new constant synthesis rate from 0 to 0.1 a.u. min^−1^ at the time of induction. In the absence of either external CSP addition or direct induction, competence is not induced (red curve). Competence development is observed either after CSP addition at *t* = 14 (blue curve) or after direct induction at *t* = 60 (orange curve). The system is refractory to CSP added 60 min after competence shut-off (pink curve). A second wave of competence is observed if, after the first addition of CSP at *t* = 14, ComX and ComW are directly induced at *t* = 60 (green curve). Blue and green curves are superimposed for the first peak of competence. (B) SsbB kinetics from normalized raw luminescence data reconstructed by applying the mathematical model of Stefan and collaborators (Stefan *et al*, 2015). Normalized experimental luminescence measurements are represented by dots. Curves were smoothed using the R function *loess* with a span parameter value of 0.2. *S. pneumoniae* R3932 is inoculated in C+Y medium at OD 550nm 0.008 (see experimental procedures). R3932 strain produces ComX and ComW following addition of either CSP or BIP. 100 ng/mL of external CSP is added at *t* = 14 and 60 min. 500 ng/mL of external BIP is added at *t*= 60. The experiment was performed three times, with similar results.

## Discussion

In this paper we report a mathematical modeling of the genetic circuit governing competence of *S. pneumoniae* that takes account of all we presently know of the system. Although competence induction and shut-off is well documented, our simulations provide mechanistic insights into their temporal regulation which we have confirmed experimentally. By mining the literature on competence regulation we developed an initial cellular scale model of CSP-induced competence regulation which underlined the incompleteness of our present knowledge on competence shut-off. Indeed, this initial model shows discrepancies between simulated and experimental data as pointed out on Figure 3 for the three different genetic backgrounds. Moreover, despite the integration of the two mechanisms known to be involved in competence shut-off, our simulation could not reproduce the competence shut-off observed in a *clpP* mutant where ComX and ComW are stable. We concluded that an unknown actor might be involved in late gene transcription termination, as proposed by Piotrowski and collaborators (Piotrowski *et al*, 2009) and Weng and collaborators (Weng *et al*, 2013). Hence, we introduced in our modeling a hypothetical actor that we named ComZ. To decipher how ComZ acts, we conceived four alternative hypotheses, while allowing ComZ to be the product of either an early or a late *com* gene.

### Competence shut-off is multifactorial

Among the eight alternative models tested (Table 3), that which best reproduces the experimental data is the one in which ComZ is the product of a late *com* gene that interacts directly with ComW to impair the formation of the active form of ComX. This model accurately recapitulates the dynamics of the interlinked ComE- and ComX-dependent transcriptional cascades. A very good agreement between the simulated and measured kinetics is obtained for the wild type strain as well as for the *dprA* and *clpP* mutants. Nevertheless, experimental time-series data are still needed for confirmation of the *clpP* simulation. Recently published data (Tovpeko *et al*, 2016) support this prediction since they demonstrate that correct shut-off is ineffective without ComW. The authors proposed that ComW may play a role in the shut-off of late gene expression, and our simulations predict that ComW is the target of an unknown late *com* gene product.

Another hypothesis concerns competition for core RNA polymerase between the active form of ComX and the housekeeping σ^A^ factor, since spontaneous suppressor mutations in a *comW*^−^ background were located in the gene *rpoD* coding for σ^A^ (Tovpeko & Morrison, 2014). In these mutants the affinity of σ^A^ for RNA polymerase could be weakened, thereby promoting σ^X^ binding. Since the transcriptome results show that *rpoD* behaves like a late *com* gene despite the absence of an upstream combox (Peterson *et al*, 2000, 2004), this hypothesis would correspond to our second best model, which posits that late ComZ competes with the active form of ComX for binding to core RNA polymerase. Even if the fit between experimental and simulated data is not as good as in the best model (compare Figures 4 and S5), especially for the wild type strain, the hypothesis of competition between σ^A^ and the active form of σ^X^ could not be excluded.

To reconcile these two observations, the most parsimonious hypothesis would be that in our two models the late ComZ corresponds to σ^A^, thus fulfilling both functions *viz.* interaction with ComW to impede the formation of the active form of σ^X^, and competition with this active form for binding to RNA polymerase. Indeed, since a tenfold increase in ComX-dependent *rpoD* expression is observed, we may assume that it will be followed by an increase of the number of σ^A^ molecules in the cell. Some of these molecules could interact with ComW for shutting off the synthesis of the active form of ComX while the others will bind to the core RNA polymerase to allow the cell to return to its original state of growth. Implementation of this hypothesis in our modeling gives an AIC value of −5900.4, slightly smaller than that obtained with our best model, and enhances slightly the fitting of experimental and simulated protein kinetics (Appendix Figure S13). However, interactions between ComW and σ^A^ need to be experimentally clarified as contradictory results have been obtained in two independent studies using yeast two-hybrid approaches. Indeed, interactions between ComW and σ^A^ have been identified by Wuchty and collaborators (Wuchty *et al*, 2017) but not by Tovpeko and collaborators (Tovpeko *et al*, 2016).

Therefore, besides the action of DprA in shutting off early *com* gene expression, our modeling predicts that another mechanism involving ComW and σ^A^ shuts off late *com* gene expression. Thus, competence shut-off appears tightly regulated at the transcriptional level both through control of the ComE master regulator of the early *com* genes and through exclusion of the σ^X^ factor required for late *com* gene expression.

### Investigation of system dynamics under specific experimental conditions

We further validated the predictive ability of the model by investigating its behavior on new aspects of the competence cycle *viz.* the effect of the basal rates of early competence gene expression on the spontaneous shift to competence, the ability for reproducing the blind-to-CSP period and the impact of the initial pH of the growth medium on competence development. We simulated the system dynamics under these specific experimental conditions and made predictions on the system components that could be involved in the observed behaviors. Experimental approaches were designed to test these predictions.

Our prediction that coordinated rates of ComCDE and ComAB basal expression are required to elicit spontaneous competence has been previously observed experimentally (Martin *et al*, 2000; Guiral *et al*, 2006). The simulated kinetics obtained for ComE, ComE∼P and SsbB reveal an almost constant value of the ratio ComE∼P/ComE (between 4e^−03^ and 6e^−03^). We propose that below this value ComE∼P cannot compete efficiently with ComE for P_*comC*_ binding and so cannot set up the positive feedback loop. At a deeper level, increased expression specifically of ComAB shortened the time taken to induce competence and conversely repression of *comAB* prolonged it (Figure 7 and Figure S12). This conclusion is supported by the behavior of mutants with raised levels of *comAB* expression (Martin *et al*, 2000; Claverys & Havarstein, 2002). The simulation results are also correlated with the loss of spontaneous competence induction in a mutant with a ten-fold increase in the basal level of *comCDE* transcription (Guiral et al 2006). Increased *comAB* gene expression in this mutant restored competence, suggesting that this phenotype was partly due to an imbalance in the positive feedback loop.

Through simulation and experimentation we demonstrate that the blind-to-CSP period is due to the presence, after competence shut-off, of residual stable DprA proteins that block the ComD/E pathway through the sequestration of ComE∼P, preventing the development of a second wave of competence.

Finally, our mathematical modeling predicts that the pH of the growth medium affects both affinity of ComD for CSP and CSP-induced ComD autophosphorylation. Indeed, by tuning concurrently the values of the two corresponding ODE parameters, our simulations can reproduce the shape of the various experimental curves. We show that signal transduction in the ComABCDE core sensing machinery detects alkaline pH through a direct effect on the binding of CSP to ComD, boosting signal transmission. Experimental results obtained for the histidine kinase AgrC from *S. aureus* have shown that the binding of the cognate ligand to one or both transmembrane domain(s) of an AgrC homodimer induces or stabilizes a conformational change in the corresponding cytoplasmic dimerization-histidine-phosphotransfer subdomains (George Cisar *et al*, 2009). Intermolecular interactions across the dimer interface induce or stabilize functionally parallel conformations in both protomers. The concerted formation of the activated conformational state in both protomers leads to the *trans*-autophosphorylation of each histidine by the contralateral kinase subdomain (George Cisar *et al*, 2009). Alkaline pH, by increasing the affinity of the ComD dimer for CSP, may favor the conformational change of the cytoplasmic domains and stabilize the active dimer conformation thus increasing the ComD autophosphorylation rate. Acidic pH, by reducing the affinity of the ComD dimer for CSP, would have the opposite effect.

### Adaptation of the model to other streptococci

While the model was built for *S. pneumoniae*, it could be adapted to other streptococcus species belonging to the *S. mitis* and *S. anginosus* groups as long as their genomes contain an ortholog of the *comW*gene. Although the *comCDE*, *comX*and *dprA* genes are conserved in all the species of these two groups, we were unable to identify orthologs of *comW* in the available genome sequences of *S. gordonii*, *S. intermedius*, *S. constellatus*, *S. sanguinis*, *S. parasanguinis*, or in *S. anginosus* with the exception of strains C1051 and J4211. For these species, it remains to be determined whether other proteins fulfil the roles of ComW in stabilizing and activating ComX or if ComX does not require any σ factor activator for binding to core RNA polymerase.. In the latter case, our model should be modified to eliminate reactions involving ComW and to allow direct synthesis of the active form of ComX through ComE∼P activation of *comX* expression. Otherwise, the model could be used with no adjustment of network topology. However, if experimental data are available, new parameter values should be estimated in order to better reflect the dynamic behavior of the network. Indeed, this dynamic may vary according to the species or even to the strains, as observed with the R800 and CP1250 strains of *S. pneumoniae* whose delay between early and late *com* gene expression and maximum gene transcription rates differ (Martin *et al*, 2013).

### Dynamic modeling at the cell population scale in S. pneumoniae

We have developed a model to account for the behavior of an individual cell in the transitory competence differentiation state in *S. pneumoniae.* The next step is to consider the cell population. Two recent articles (Prudhomme *et al*, 2016; Moreno-Gámez *et al*, 2017) focused on spontaneous competence development at the population scale but interpreted the results obtained differently. Moreno and collaborators (Moreno-Gámez *et al*, 2017) based their conclusions on classic quorum sensing by the free release of CSP, which leads to a synchronization of the shift to competence in the whole population. On the other hand, Prudhomme and collaborators (Prudhomme *et al*, 2016) favor the initiation of competence in a small fraction of the population that then propagates competence among the non-competent cells by distributing CSP.

The present work will allow, at a minimum, the testing of both hypotheses by its integration into more complex modeling. Approaches like agent-based or individual-based models (ABMs, IBMs) (Gorochowski *et al*, 2012; Hellweger *et al*, 2016) could be used. These models consider populations of autonomous agents, each following a set of internal rules and interacting with each other within a shared virtual environment. Our model can be used to design each agent and its evolution (competence state shift) over time depending on the environment inputs like CSP capture through contact with a competent cell and CSP free diffusion. Interactions between agents, here the cells, can also be defined differently according to the life style of the bacteria as either planktonic or in biofilm. While in the synchronization model (Moreno-Gámez *et al*, 2017) the cell population is homogeneous, in the competence propagation model (Prudhomme *et al*, 2016) the design of a dynamic model requires taking into account the non-homogeneity of the population. The population should contain at least three types of cell: competence-initiator cells that will develop spontaneous competence, CSP-induced competent cells whose competence development relies on CSP transmission, and non-competent cells that will not respond to CSP. If our present cellular model reflects CSP-induced competent cell behavior, few adjustments will be required to simulate the behavior of the other two cell types. Indeed, for competence-initiator cells we have already shown that spontaneous competence can be monitored by tuning the basal expression rates of ComCDE and ComAB. Agent-based or individual-based models have already been applied to model different biological processes in microbial populations and open-source platforms have been developed, such as INDISIM, to simulate the growth and behavior of bacterial colonies (Ginovart *et al*, 2002) - BSim for gene regulation (Gorochowski *et al*, 2012), Framework for biofilm models (Xavier *et al*, 2005) or AgentCell for chemotaxis signaling (Emonet & Cluzel, 2008).

Besides its use in the development of a population-scaled model for dynamic study of the competence circuit in *S. pneumoniae*, our model can be exploited to model more complex biological processes, such as the cell cycle, where it could be embedded as a module interconnected with other parts of the network.

## Materials and Methods

### Petri net modeling

We implement our Petri net model using Snoopy’s framework (Heiner *et al*, 2012; Marwan *et al*, 2012). We used the software Charlie (Heiner *et al*, 2015) to compute the structural invariants (T- and P-invariants) of the network. A brief, basic introduction to Petri net and structural invariants calculation is available as supplementary material (Appendix Supplementary Methods). More details on Petri net can be found in (Chaouiya, 2007) and (Koch & Heiner, 2008).

### Estimations of protein synthesis rates and protein concentration kinetics

Promoter activities were obtained by transformation of the luminescence signal following the formula proposed by Stefan and collaborators (Stefan *et al*, 2015):

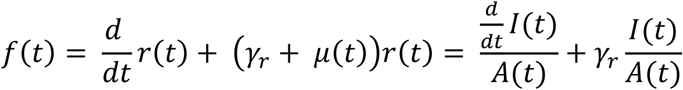

where *A*(*t*) corresponds to the background corrected absorbance value, *I*(*t*) to the luminescence intensity (in relative luminescence units; RLU) over time, *r*(*t*)=*I*(*t*)/*A*(*t*) to the quantity of luminescence per cell as a function of time (de Jong *et al*, 2010), *γ_r_* [min ^−1^] to the degradation constant of the luciferase protein and μ(*t*) [min ^−1^] to the growth rate of the bacteria. The half-life time of the luciferase used in our experiments was determined to be 15 min (Prudhomme & Claverys, 2007), leading to *μ_r_*= 0.04621. The reporter signal is expressed in RLU and the promoter activity in RLU min^−1^.

The rate of change in concentration over time of the protein of interest *p*(*t*) [RFU min^−1^] was calculated using the following equation (Stefan *et al*, 2015), which corrects for the difference in half-life between the reporter luciferase and the protein whose gene activity is measured :

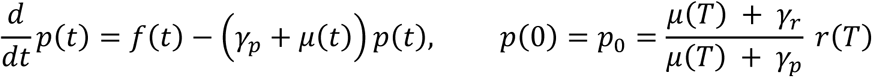

where *γ_p_*[min^−1^] is the degradation constant of the protein and *μ(T)* is the growth rate of bacteria at the end of the preculture procedure (at time T). Based on published results and for simplicity, we considered only two different life times, one for ComD, ComE, DprA and ComAB, estimated at 80 min (*t*_1/2_= 55.45 min andγ_*p*_ = 0.0125) and the other for ComX, ComW and SsbB, estimated at 8 min (*t*_1/2_= 5.45 min and γ_*P*_ = 0.125) ((Piotrowski *et al*, 2009) for ComX and ComW; (Mirouze *et al*, 2013) for DprA and SsbB; (Martin *et al*, 2013) for ComE and ComD). We assume that competence gene expression is at steady-state at the beginning of the experiment, so *μ*(*T*) = 0 and the initial protein concentration *p0* depends only on the protein and luciferase degradation constants and on the ratio *r*(T).

Equations were solved by numerical integration using the Euler method implemented in R software from the deSolve package. Finally, the protein concentrations obtained were normalized between 0 and 1 with respect to maximum protein concentration values obtained over all computed data sets.

### Model parameter estimation

The parameter inference problem for ordinary differential equation models is usually formulated as an optimization problem with an objective function that has to be minimized by adjusting the values of the model parameters. A common choice for computing this objective function is to calculate the sum of squared errors between measurements and model predictions. In COPASI, for a set of parameters P, the objective function is given by the following formula (Hoops *et al*, 2006):

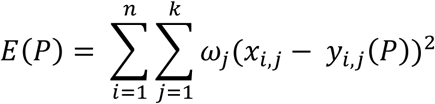

Where *n* is the number of data point, *k* is the number of variables, *y_i_*,(P)are the simulated data corresponding to the experimental data *X*_*i*,*j*_ and *ω_j_* is a weight that gives a similar importance in the fit to all trajectories of each variable. To calculate the weights, we used the mean square calculation method.

The particle swarm algorithm optimization method (PSO) implemented in COPASI (Hoops *et al*, 2006) was used to estimate the parameter values. Details on PSO can be found in (Eberhart & Kennedy, 1995) and in (Poli *et al*, 2007). We have bounded the parameter search space by using the constraints set up on the different parameters of the model that are described in the result section of the manuscript. We run the PSO algorithm by using the default parameters proposed in COPASI, except for the number of iterations, which was increased from 2000 to 10000.

Simulations of the network behavior were performed using LSODA deterministic solver (Petzold, 1983) for ODE numerical integration. LSODA automatically determines if a system of ordinary differential equations can be solved more efficiently by a class of methods suited for non-stiff problems or by a class of methods dedicated to stiff problems. LSODA was run with the default parameters implemented in COPASI.

### pH effect on competence development

Neither the *comA* mutant R1205 (Martin *et al*, 2013), which cannot export CSP, nor the comC mutant R1694 (Martin *et al*, 2010), which cannot synthesize it, develop competence naturally, but both do so upon addition of synthetic CSP. To observe competence development as a function of CSP concentration and culture medium pH we inoculated C+Y medium (Tomasz & Hotchkiss, 1964) at pH 7.27 with each strain at OD_550_ 0.01 and incubated the cultures at 37°C until the OD reached 0.12. The cells were then washed by centrifugation/resuspension in the same medium, concentrated to OD 1, and kept on ice. The cells were diluted to OD ∼0.03 in pre-warmed (37°C) C+Y medium adjusted to the desired pH using NaOH or HCl containing luciferin, as previously described (Prudhomme & Claverys, 2007). The diluted cells (300 μl of aliquots) were immediately transferred to clear bottomed wells of a 96-well white NBS micro plate (Corning) at 37°C. CSP was then added at the desired concentration. Relative luminescence units (RLU) values were recorded every minute throughout incubation at 37°C in a Varioskan Flash (Thermo 399 Electron Corporation) luminometer. The pH in replicate experiments varied slightly from the nominal values but these variations did not significantly affect the competence induction profiles.

### Effect of DprA-mediated blocking of the ComD/Epathway on the blind-to-CSP period

Strain R3932 was constructed as follows: i) strand overlap extension using the primers MB26, −27, −28, −29 to substitute comC2 for comC1 in the strain R1036 by the Janus methods described in (Sung *et al*, 2001); ii) The strain was then converted to the wild type *rpsL* locus by transformation (R3369); iii) introduction of *ssb::luc* (Martin *et al*, 2000) by transformation to yield R3584; iv) cloning of a *comX1*-*comW* fragment created by strand-overlap extension using primers MB54,−56,−57,−58 between the BamHI and Nco1 sites of pCEPR-luc (Johnston *et al*, 2016), and transformation R3584 by the resulting plasmid to yield R3932 (*comC2D1*, *ssbB::luc (Cm)*, *pcepR*-*comX1*-*comW(kanR))*. Competence was monitored as above, using the *ssbB::luc* transcriptional fusion. Inducing peptides were added at final concentrations of 100 ng/mL (CSP) or 500 ng/mL (BIP). Primer sequences are listed in Appendix Table S2.

### Data availability

The model was deposited in BioModels (Chelliah *et al*, 2015) and assigned the identifier MODEL1803300001. The datasets used and/or analyzed during the current study are available from the corresponding author on request.

## Acknowledgements

We thank Yves Quentin for its help with analyses of data from growth medium pH value experiments and for useful discussions during manuscript preparation. We thank Mike Chandler and Dave Lane for critical reading of the manuscript. This work was supported by institutional grants from the CNRS (Centre National de la Recherche Scientifique). MW was supported by an MESR (Ministère de l’Enseignement Supérieur et de la Recherche) fellowship.

## Author contributions

Conceived and designed the mathematical modeling: GF MW. Conceived and designed the experiments: MP MB. Performed the theoretical experiments: MW GF. Performed the biological experiments: MP MB. Analyzed the data: GF MW MP MB PP. Wrote the paper: GF MP. All authors read and approved the final manuscript.

